# Multiple bumps can enhance robustness to noise in continuous attractor networks

**DOI:** 10.1101/2022.02.22.481545

**Authors:** Raymond Wang, Louis Kang

## Abstract

A central function of continuous attractor networks is encoding coordinates and accurately updating their values through path integration. To do so, these networks produce localized bumps of activity that move coherently in response to velocity inputs. In the brain, continuous attractors are believed to underlie grid cells and head direction cells, which maintain periodic representations of position and orientation, respectively. These representations can be achieved with any number of activity bumps, and the consequences of having more or fewer bumps are unclear. We address this knowledge gap by constructing 1D ring attractor networks with different bump numbers and characterizing their responses to three types of noise: fluctuating inputs, spiking noise, and deviations in connectivity away from ideal attractor configurations. Across all three types, networks with more bumps experience less noise-driven deviations in bump motion. This translates to more robust encodings of linear coordinates, like position, assuming that each neuron represents a fixed length no matter the bump number. Alternatively, we consider encoding a circular coordinate, like orientation, such that the network distance between adjacent bumps always maps onto 360 degrees. Under this mapping, bump number does not significantly affect the amount of error in the coordinate readout. Our simulation results are intuitively explained and quantitatively matched by a unified theory for path integration and noise in multi-bump networks. Thus, to suppress the effects of biologically relevant noise, continuous attractor networks can employ more bumps when encoding linear coordinates; this advantage disappears when encoding circular coordinates. Our findings provide motivation for multiple bumps in the mammalian grid network.

## Introduction

Continuous attractor networks (CANs) sustain a set of activity patterns that can be smoothly morphed from one to another along a low-dimensional manifold (Amari, 1977; Ermentrout and Cowan, 1979; Milnor, 1985). Network activity is typically localized into attractor bumps, whose positions along the manifold can represent the value of a continuous variable. These positions can be set by external stimuli, and their persistence serves as a memory of the stimulus value. Certain CAN architectures are also capable of a feature called path integration. Instead of receiving the stimulus value directly, the network receives its changes and integrates over them by synchronously moving the attractor bump (Cannon et al., 1983; McNaughton et al., 1991; Seung, 1996). Path integration allows systems to estimate an external state based on internally perceived changes, which is useful in the absence of ground truth.

Path-integrating CANs have been proposed as a mechanism through which brains encode various physical coordinates. Head direction cells in mammals and compass neurons in insects encode spatial orientation by preferentially firing when the animal faces a particular direction relative to landmarks (Fig. 1A, top; Taube et al., 1990; Seelig and Jayaraman, 2015). They achieve this as members of 1D CANs whose attractor manifolds have ring topologies (Skaggs et al., 1995; Zhang, 1996). For the case of compass neurons, a ring structure also exists anatomically, and its demonstration of continuous attractor dynamics is well-established (Seelig and Jayaraman, 2015; Kim et al., 2017; Turner-Evans et al., 2017; Green et al., 2017). Grid cells in mammals encode position by preferentially firing at locations that form a triangular lattice in 2D space (1D analogue in Fig. 1A, bottom; Hafting et al., 2005). They are thought to form a 2D CAN with toroidal topology (McNaughton et al., 2006; Fuhs and Touretzky, 2006; Guanella et al., 2007; Burak and Fiete, 2009), and mounting experimental evidence supports this theory (Yoon et al., 2013; Gu et al., 2018; Gardner et al., 2019, 2022). The ability for head direction cells, compass neurons, and grid cells to maintain their tunings in darkness without external cues demonstrates that these CANs can path integrate (Goodridge et al., 1998; Seelig and Jayaraman, 2015; Hafting et al., 2005).

**Figure 1:**
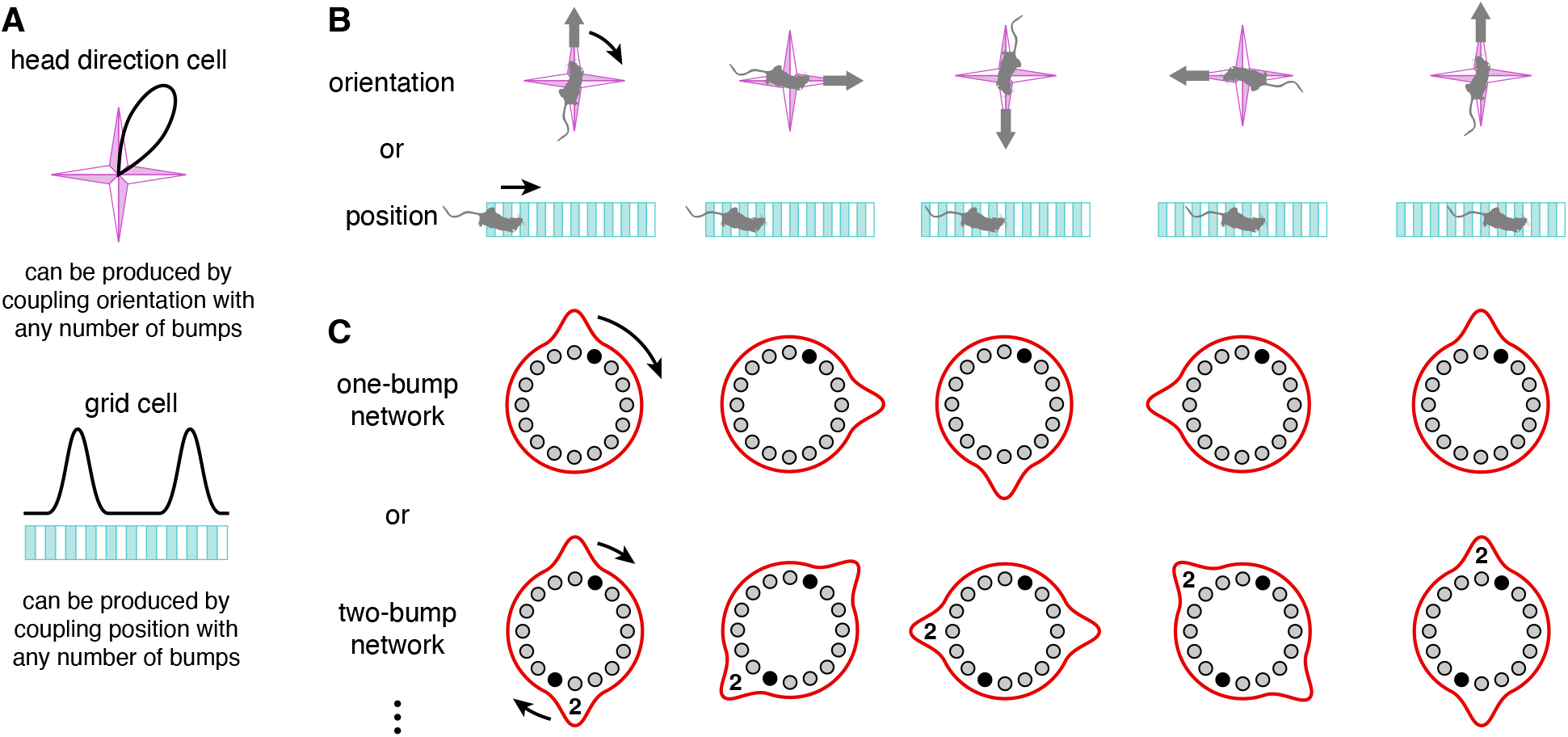
Continuous attractor networks with any number of bumps can produce head direction cells and grid cells. (**A**) Desired tuning curves of a head direction cell and a 1D grid cell. (**B**) Orientation and position coordinates whose changes drive bump motion. (**C**) One- and two-bump ring attractor networks. Each black neuron produces the desired tuning curves in **A**. In the two-bump network, the coupling to coordinate changes is half as strong, and the second bump is labeled for clarity.

CANs also appear in studies of other brain regions and neural populations. Signatures of continuous attractor dynamics have been detected in the prefrontal cortex during spatial working memory tasks (Constantinidis and Wang, 2004; Edin et al., 2009; Wimmer et al., 2014). Theorists have further invoked CANs to explain place cells (Tsodyks and Sejnowski, 1995; Samsonovich and McNaughton, 1997), hippocampal view cells (Stringer et al., 2005), eye tracking (Cannon et al., 1983; Seung, 1996), visual orientation tuning (Ben-Yishai et al., 1995; Somers et al., 1995), and perceptual decision making (Brody et al., 2003; Machens et al., 2005). Thus, CANs are a crucial circuit motif throughout the brain, and better understanding their performance would provide meaningful insights into neural computation.

One factor that strongly affects the performance of CANs in path integration is biological noise. To accurately represent physical coordinates, attractor bumps must move in precise synchrony with the animal’s trajectory. Hence, the bump velocity must remain proportional to the driving input that represents coordinate changes (Burak and Fiete, 2009). Different sources of noise produce different types of deviations from this exact relationship, all of which lead to path integration errors. While noisy path-integrating CANs have been previously studied (Zhang, 1996; Stringer et al., 2002; Wu et al., 2008; Burak and Fiete, 2009), these works did not investigate of role of bump number. CANs with different connectivities can produce different numbers of attractor bumps, which are equally spaced throughout the network and perform path integration by moving in unison (Stringer et al., 2004; Fuhs and Touretzky, 2006; Burak and Fiete, 2009). Two networks with different bump numbers have the same representational capability (Fig. 1). They can share the same attractor manifold and produce neurons with identical tuning curves, as long as the coupling strength between bump motion and driving input scales appropriately. The computational advantages of having more or fewer bumps are unknown.

Our aim is to elucidate the relationship between bump number and robustness to noise. We first develop a rigorous theoretical framework for studying 1D CANs that path integrate and contain multiple bumps. Our theory predicts the number, shape, and speed of bumps. We then introduce three forms of noise. The first is Gaussian noise added to the total synaptic input, which can represent fluctuations in a broad range of cellular processes occurring at short timescales. The second is Poisson spiking noise. The third is noise in synaptic connectivity strengths; the ability for bumps to respond readily to driving inputs is generally conferred by a precise network architecture. We add Gaussian noise to the ideal connectivity and evaluate path integration in this setting. The first two forms of noise are independent over time and neurons, in contrast to the third. We find that networks with more bumps can better resist all three forms of noise under certain encoding assumptions. These observations are explained by our theoretical framework with simple scaling arguments. The following Results section presents all simulation findings and major theoretical conclusions; complete theoretical derivations are found in the Theoretical model section.

## Results

### Bump formation in a ring attractor network

We study a 1D ring attractor network that extends the model of Xie et al. (2002) to allow for multiple attractor bumps. It contains two neural populations *α ∈ {*L, R*}* at each network position *x*, with *N* neurons in each population (Fig. 2A). Each neuron is described by its total synaptic input *g* that obeys the following dynamics:

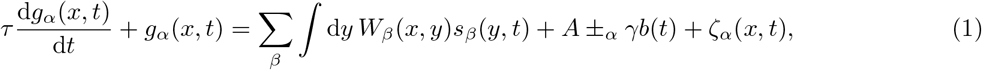

where *±*_L_ means *−* and *±*_R_ means +. Aside from spiking simulations, firing rates *s* are given by

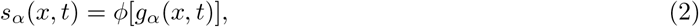

where *ϕ* is a nonlinear activation function. For all simulations in this Results section, we take *ϕ* to be the rectified linear unit (ReLU) activation function (Eq. 35). Our theoretical formulas for diffusion coefficients and velocities in this section also assume a ReLU *ϕ*. In the Appendix, we consider a logistic *ϕ* instead and find that all major conclusions are preserved (Fig. 9), and in the Theoretical methods section, we derive most expressions for general *φ. W* is the synaptic connectivity and only depends on the presynaptic population *β*. It obeys a standard continuous attractor architecture based on local inhibition that is strongest at an inhibition distance *l*. Each population has its synaptic outputs shifted by a small distance *ξ* ≪ *l* in opposite directions. We use the connectivity profile described in Fig. 2B and Eq. 38 for all simulations, but all theoretical expressions in this Results section are valid for any *W*. *A* is the resting input to all neurons. The driving input, or drive, *b* is proportional to changes in the coordinate encoded by the network; for the physical coordinates in Fig. 1B, it represents the animal’s velocity obtained from self-motion cues. In our results, *b* is constant in time. It is coupled to the network with strength *γ*. We will consider various forms of noise *ζ*. Finally, *τ* is the neural time constant.

**Figure 2:**
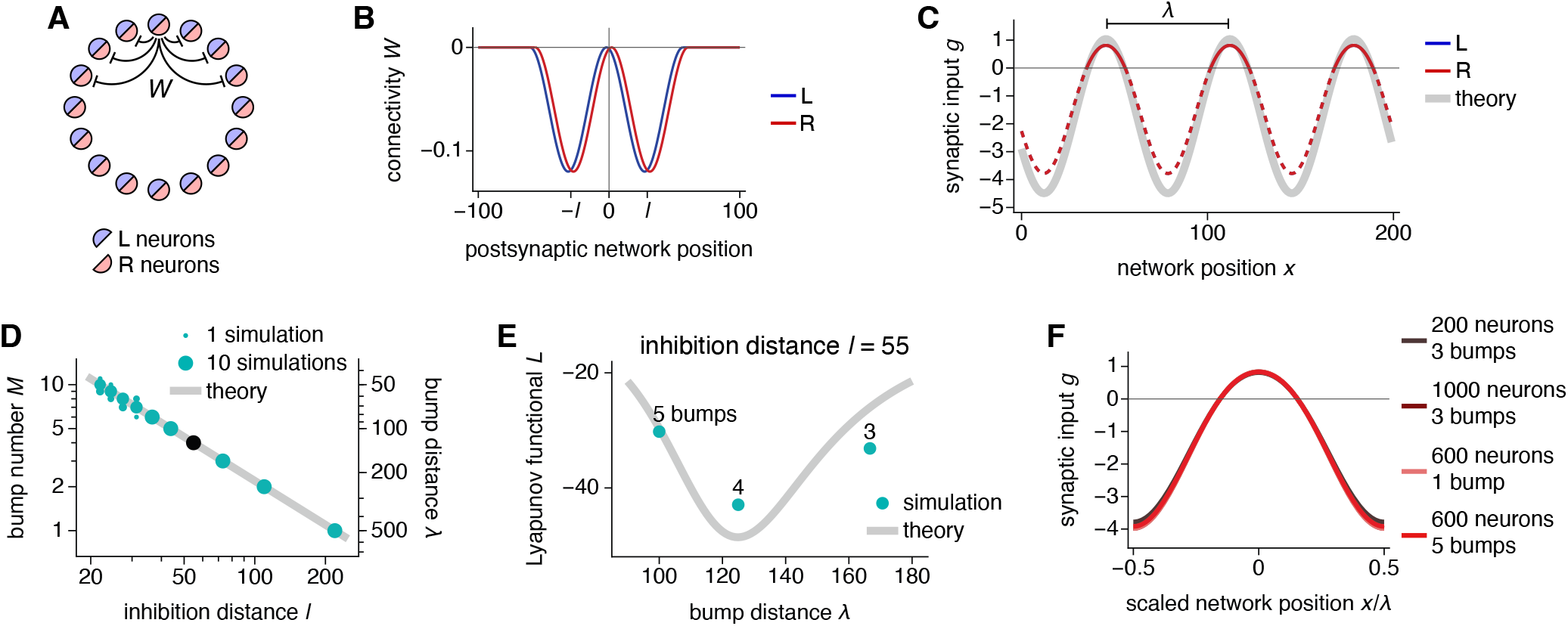
Bump formation in a ring attractor network. (**A**) Network schematic with populations L and R and locally inhibitory connectivity *W*. (**B, C**) Networks with 200 neurons and 3 bumps. (**B**) Connectivity weights for a neuron at the origin. The inhibition distance is *l* = 29 and the connectivity shift is *ξ* = 2. (**C**) Steady-state synaptic inputs. Curves for both populations lie over each other. With a ReLU activation function, the firing rates follow the solid portions of the colored lines and are 0 over the dashed portions. The bump distance is *λ* = 200*/*3. Thick gray line indicates Eq. 4. (**D, E**) Networks with 500 neurons. (**D**) More bumps and shorter bump distances are produced by smaller inhibition distances. Points indicate data from 10 replicate simulations. Line indicates Eq. 5. (**E**) The inhibition distance *l* = 55 corresponds to the black point in **D** with *λ* = 125 and *M* = 4. These values also minimize the Lyapunov functional (Eq. 6), which varies smoothly across *λ* for infinite networks (line) and takes discrete values for finite networks (points). (**F**) The scaled bump shape remains invariant across network sizes and bump numbers, accomplished by rescaling connectivity strengths according to Eq. 7. Curves for different parameters lie over one another.

With no drive *b* = 0 and no noise *ζ* = 0, the network dynamics in Eqs. 1 and 2 can be simplified to

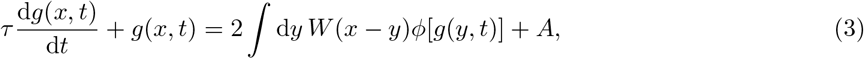

where 2*W* (*x − y*) = Σ_*β*_ *W*_*β*_(*x, y*) and the synaptic inputs *g* are equal between the two populations. This baseline equation evolves towards a periodic steady-state *g* with approximate form (see also Widloski, 2015)

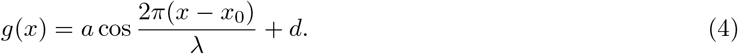

Expressions for *a* and *d* are given in the Theoretical model section (Eq. 60). The firing rates *s*(*x*) = *ϕ*[*g*(*x*)] exhibit attractor bumps with periodicity *λ*, a free parameter that we call the bump distance (Fig. 2C). *x*^0^ is the arbitrary position of one of the bumps. It parameterizes the attractor manifold with each value corresponding to a different attractor state up to *λ*.

The bump number *M* = *N/λ* is determined through *λ*. It can be predicted by the fastest-growing mode in a linearized version of the dynamics (Eq. 43; Sorscher et al., 2019; Khona et al., 2022). The mode with wavenumber *q* and corresponding wavelength 2*π/q* grows at rate 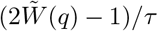 where 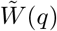 is the Fourier transform of *W* (*x*). Thus,

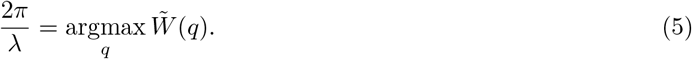

Figure 2D shows that simulations follow the predicted *λ* and *M* over various inhibition distances *l*. Occasionally for small *l*, a different mode with a slightly different wavelength will grow quickly enough to dominate the network. A periodic network enforces an integer bump number, which discretizes the allowable wavelengths and prevents changes in *λ* and *M* once they are established. In an aperiodic or infinite system, the wavelength can smoothly vary from an initial value to a preferred length over the course of a simulation (Burak and Fiete, 2009; Kang and Balasubramanian, 2019). To determine this preferred *λ* theoretically, we notice that the nonlinear dynamics in Eq. 3 obey the Lyapunov functional

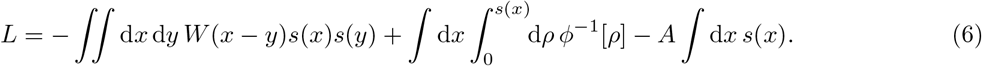

In the Theoretical model section, we find for ReLU *ϕ* that *L* is minimized when *q* = 2*π/λ* maximizes 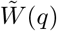 (Eq. 66). This is the same condition as for the fastest-growing mode in Eq. 5 (Fig. 2E). In other words, the wavelength *λ* most likely to be established in a periodic network is the preferred bump distance in an aperiodic or infinite system, up to a difference of one fewer or extra bump due to discretization.

We now understand how to produce different bump numbers *M* in networks of different sizes *N* by adjusting the inhibition distance *l*. To compare networks across different values of *M* and *N*, we scale the connectivity strength *W* according to

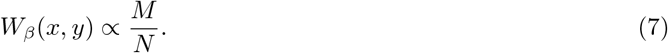

This keeps the total connectivity strength per neuron ∫ d*x W*_*β*_(*x, y*) constant over *M* and *N*. In doing so, the shape of each attractor bump as a function of scaled network position *x/λ* remains invariant (Fig. 2F). Thus, Eq. 7 isolates our comparisons across *M* and *N* to those variables themselves and removes any influence of bump shape. In the Appendix, we consider the alternative without this scaling and find that many major results are preserved (Fig. 10).

### Bump dynamics: path integration and diffusion

The drive *b* produces coherent bump motion by creating an imbalance between the two neural populations. A positive *b* increases input to the R population and decreases input to the L population (Fig. 3A). Because the synaptic outputs of the former are shifted to the right, the bump moves in that direction. Similarly, a negative *b* produces leftward bump motion. The bump velocity *v*^drive^ can be calculated in terms of the baseline firing rates *s*(*x*) obtained without drive and noise (see also Xie et al., 2002; Mosheiff and Burak, 2019):

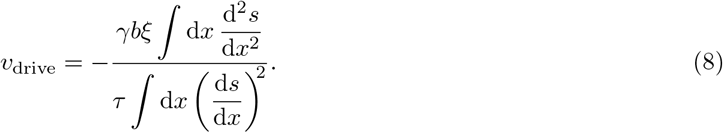

As a note, these integrals, as well as subsequent ones, do not include the singular points at the edges of attractor bumps. Equation 8 states that bump velocity is proportional to drive *b* and connectivity shift *ξ*, which is reflected in our simulations, with some deviation at larger *ξ* (Fig. 3B, C). The strict proportionality between *v* and *b* is crucial because it implies faithful path integration (Burak and Fiete, 2009). If *b*(*t*) represents coordinate changes (such as angular or linear velocity in Fig. 1B), then the bump position *θ*(*t*) will accurately track the coordinate itself (orientation or position).

**Figure 3:**
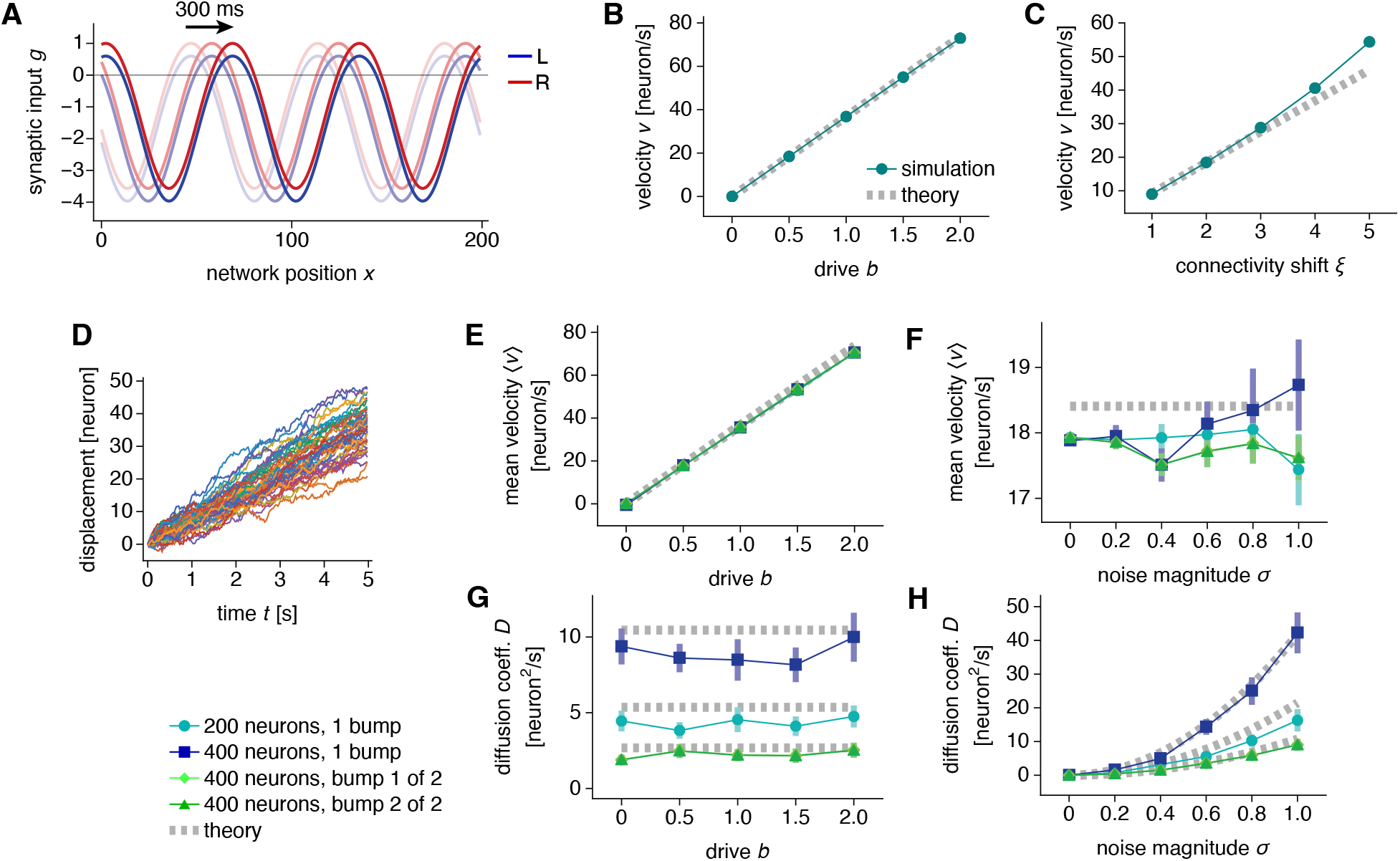
Dynamics in a ring attractor network. (**A–C**) Networks with 200 neurons and 3 bumps. (**A**) Synaptic inputs for populations L and R under drive *b* = 2. Snapshots taken at 150 ms intervals demonstrate rightward motion. (**B**) Bump velocity is proportional to drive. The connectivity shift is *ξ* = 2. (**C**) Bump velocity is largely proportional to connectivity shift. The drive is *b* = 0.5. (**D–H**) Networks with synaptic input noise. (**D**) Bump displacements for 48 replicate simulations demonstrating diffusion with respect to coherent motion. Networks with 200 neurons and 1 bump. (**E, F**) Mean bump velocity is proportional to drive and remains largely independent of network size, bump number, and noise magnitude. (**G, H**) Bump diffusion coefficient scales quadratically with noise magnitude, remains largely independent of drive, and varies with network size and bump number. The noise magnitude is *σ* = 0.5 in **D, E**, and **G**, and the drive is *b* = 0.5 in **D, F**, and **H**. Values for both bumps in two-bump networks lie over each other. Points indicate data from 48 replicate simulations and bars indicate bootstrapped standard deviations. Dotted gray lines indicate Eqs. 8 and 10.

In contrast to drive, uncorrelated noise *ζ* produces bump diffusion. To illustrate this effect, we introduce one form of *ζ* that we call synaptic input noise. Suppose *ζ* is independently sampled for each neuron at each simulation timestep from a Gaussian distribution with mean 0 and variance *σ*^2^. Loosely, it can arise from applying the central limit theorem to the multitude of noisy synaptic and cellular processes occurring at short timescales. Then,

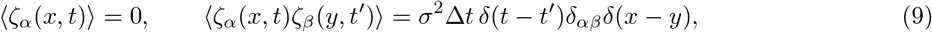

where the timestep Δ*t* sets the resampling rate of *ζ*, and angle brackets indicate averaging over an ensemble of replicate simulations. Input noise causes bumps to diffuse away from the coherent driven motion (Fig. 3D).

The mean velocity ⟨*v*⟩ remains proportional to drive *b*, which means that the network still path integrates on average (Fig. 3E). Since ⟨*v*⟩ is largely independent of noise magnitude *σ*, and the bump diffusion coefficient *D* is largely independent of *b*, drive and input noise do not significantly interact within the explored parameter range (Fig. 3F, G). *D* can be calculated in terms of the baseline firing rates (see also Wu et al., 2008; Burak and Fiete, 2012):

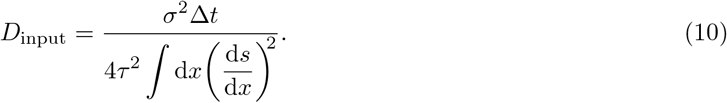

The quadratic dependence of *D* on *σ* is confirmed by simulation (Fig. 3H).

We now turn our attention to bump number *M* and network size *N*. The mean bump velocity ⟨*v*⟩ is independent of these parameters (Fig. 3E, F), which can be understood theoretically. Bump shapes across *M* and *N* are simple rescalings of one another (Fig. 2F), so derivatives of *s* with respect to *x* are simply proportional to *M* (more bumps imply faster changes) and inversely proportional to *N* (larger networks imply slower changes). Similarly, integrals of expressions containing *s* over *x* are simply proportional to *N*. In summary,

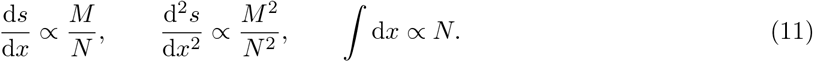

Applying these scalings to Eq. 8, we indeed expect *v*_drive_ to be independent of *M* and *N*. In contrast, Fig. 3G, H reveals that the diffusion coefficient *D* varies with these parameters. When a one-bump network is increased in size from 200 to 400 neurons, *D* increases as well, which implies greater path integration errors. This undesired effect can be counteracted by increasing the bump number from 1 to 2, which lowers *D* below that of the one-bump network with 200 neurons. These initial results suggest that bump number and network size are important factors in determining a CAN’s resilience to noise. We will explore this idea in greater detail.

### Mapping network coordinates onto physical coordinates

Before further comparing networks with different bump numbers *M* and sizes *N*, we should scrutinize the relationship between bump motion and the physical coordinate encoded by the network. After all, the latter is typically more important in biological settings. First, we consider the trivial case in which each neuron represents a fixed physical interval across all *M* and *N*; this is equivalent to using network coordinates without a physical mapping (Fig. 4A). It is suited for encoding linear variables like position that lack intrinsic periodicity, so larger networks can encode wider coordinate ranges. However, with more bumps or fewer neurons, the range over which the network can uniquely encode different coordinates is shortened. We assume that ambiguity among coordinates encoded by each bump can be resolved by additional cues, such as local features, that identify the true value among the possibilities (O’Keefe and Burgess, 2005; Sreenivasan and Fiete, 2011; Stemmler et al., 2015); this process will be examined in detail below. We leave quantities with dimensions of network distance in natural units of neurons.

**Figure 4:**
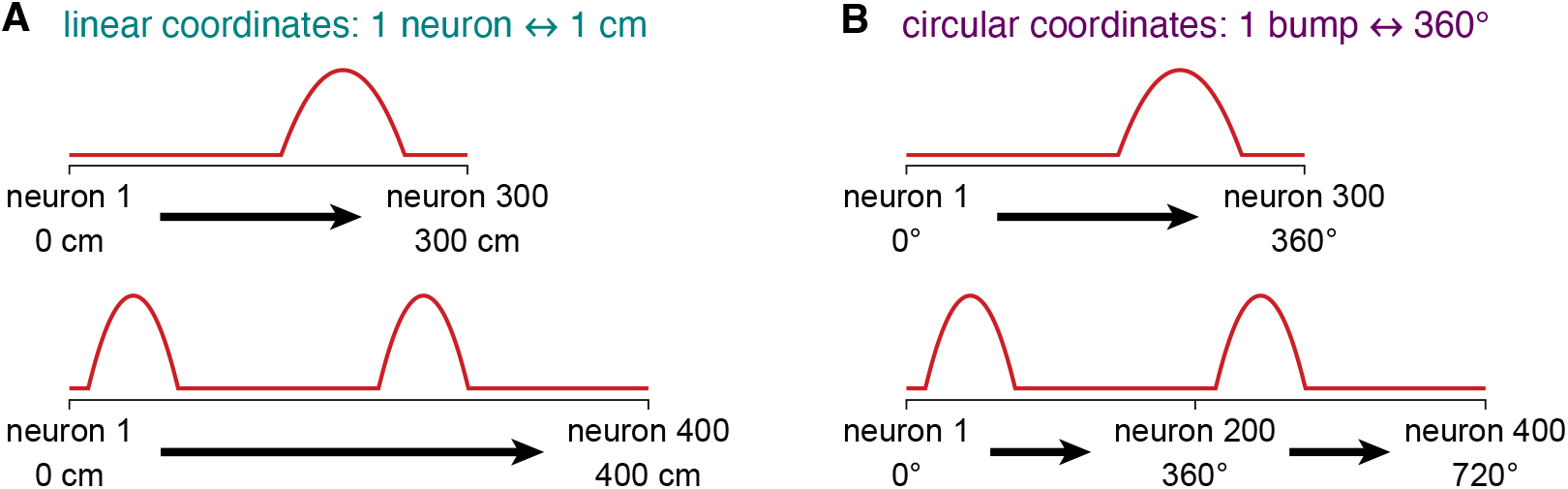
Possible mappings between network coordinates and two types of physical coordinates. (**A**) In networks encoding linear coordinates such as position, one neuron always represents a fixed physical interval. This mapping is trivial and identical to using network coordinates. (**B**) In networks encoding circular coordinates such as orientation, the bump distance always represents 360.

Multi-bump networks are intrinsically periodic, especially those with a ring architecture. A natural way for them to encode a circular coordinate like orientation would be to match network and physical periodicities. For example, the bump distance may always represent 360 across different *M* and *N* so that neurons always exhibit unimodal tuning (Fig. 4B). This relationship implies that quantities with dimensions of network distance should be multiplied by powers of the conversion factor

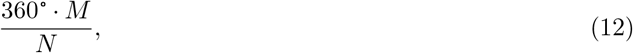

which converts units of neurons to degrees.

For circular mapping, we must also ensure that networks with different bump numbers *M* and sizes *N* path integrate consistently with one another. The same drive *b* should produce the same bump velocity *v* in units of degree/s. To do so, we rescale the coupling strength *γ* only under circular mapping:

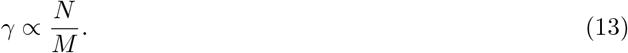

This effectively compensates for the factor of *M/N* in Eq. 12. To see this explicitly, recall that *v*_drive_ does not depend on *M* and *N* in units of neuron/s, as shown in Fig. 3E, F and previously explained through scaling arguments. Under circular mapping, *v*_drive_ would be multiplied by one power of the conversion factor in Eq. 12. Since its formula contains *γ* in the numerator (Eq. 8), *v*_drive_ receives an additional power of the rescaling factor in Eq. 13. The two factors cancel each other, so *v*_drive_ does not depend on *M* and *N* under either mapping:

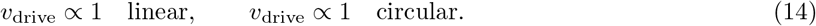

Thus, a consistent relationship between *b* and *v*_drive_ is preserved in units of both neurons/s and degrees/s.

Of course, there are other possible mappings between network and physical coordinates across bump numbers and network sizes, but for the rest of our paper, we will consider these two. To be clear, networks with the same ring architecture are used for both linear and circular mappings. We will see how noise affects encoding quality in either case.

### More bumps improve robustness to input and spiking noise under linear mapping

We now revisit the effect of input noise on bump diffusion, as explored in Fig. 3D–H. We measure how the diffusion coefficient *D* varies with bump number *M* and network size *N* under linear and circular mappings. Under linear mapping, *D* decreases as a function of *M* but increases as a function of *N* (Fig. 5A, B). Thus, more bumps attenuate diffusion produced by input noise, which is especially prominent in large networks.

**Figure 5:**
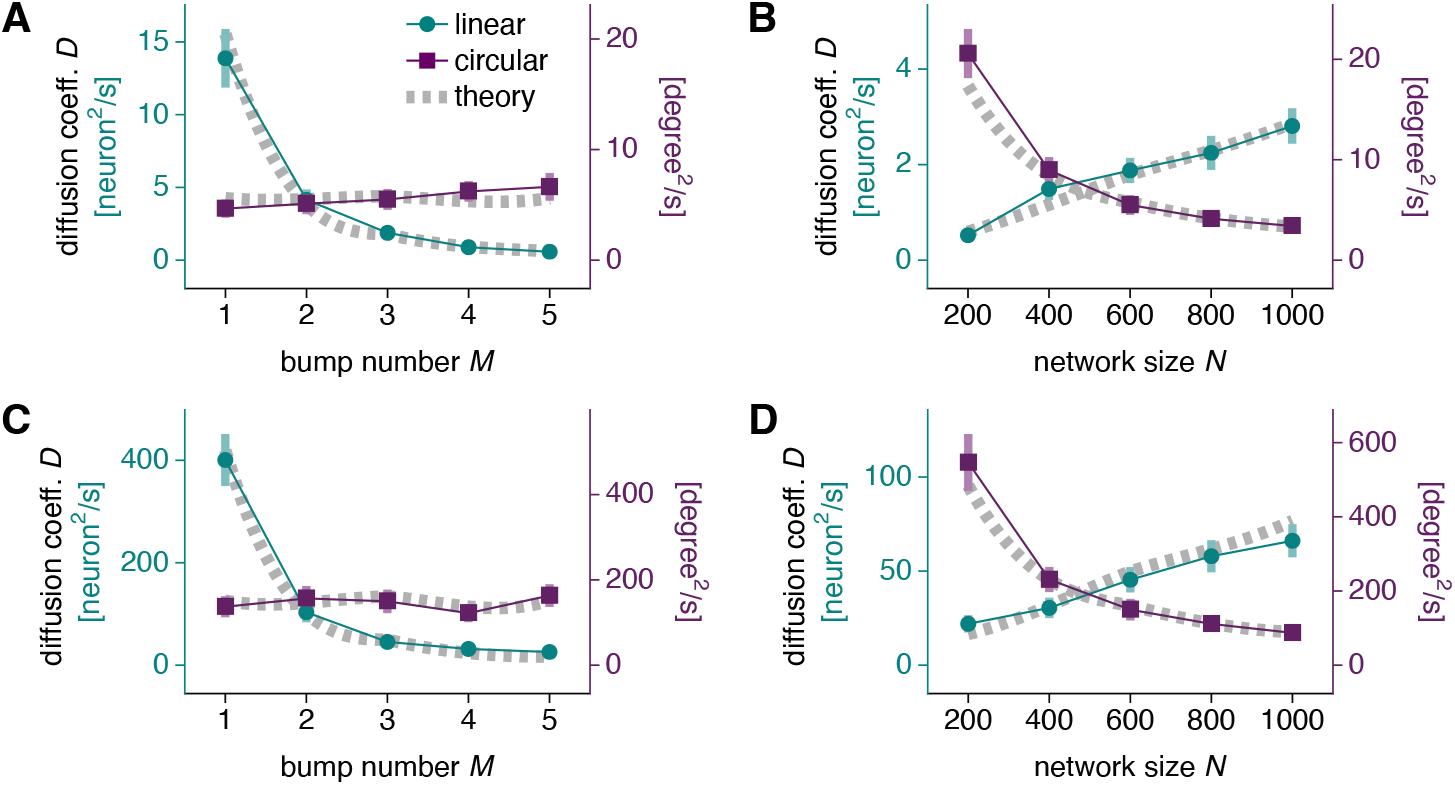
Bump diffusion due to input and spiking noise. (**A, B**) Networks with synaptic input noise of magnitude *σ* = 0.5 and drive *b* = 0.5. Dotted gray lines indicate Eq. 10. (**A**) Diffusion decreases with bump number under linear mapping and remains largely constant under circular mapping. Networks with 600 neurons. (**B**) Diffusion increases with network size under linear mapping and decreases under circular mapping. Networks with 3 bumps. (**C, D**) Same as **A, B**, but for networks with Poisson spiking noise instead of input noise. Dotted gray lines indicate Eq. 20. Points indicate data from 48 replicate simulations and bars indicate bootstrapped standard deviations.

However, for circular coordinates, *D* remains largely constant with respect to *M* and decreases with respect to *N* (Fig. 5A, B). Increasing the number of bumps provides no benefit. These results can be understood through Eqs. 10, 11, and 12, which predict

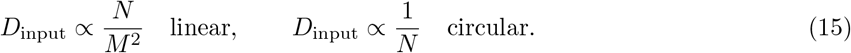

Two powers of the conversion factor in Eq. 12 account for the differences between the two mappings.

Next, we investigate networks with spiking noise instead of input noise. To do so, we replace the deterministic formula for firing rate in Eq. 2 with

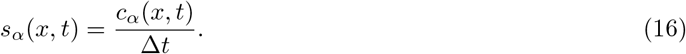

Here, *s* is a stochastic, instantaneous firing rate given by the number of spikes *c* emitted in a simulation timestep divided by the timestep duration Δ*t*. We take the *c*’s to be independent Poisson random variables driven by the deterministic firing rate:

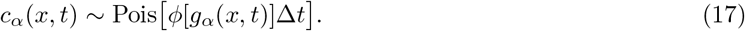

As fully explained in the Theoretical model section (Eq. 99), we can approximate this spiking process by the rate-based dynamics in Eqs. 1 and 2 with the noise term

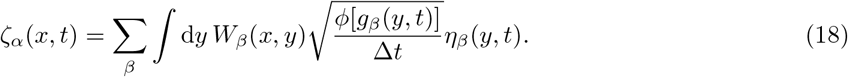

The *η*’s are independent random variables with zero mean and unit variance:

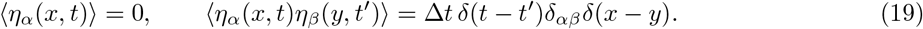

As for Eq. 9, the simulation timestep Δ*t* sets the rate at which *η* is resampled. This spiking noise produces bump diffusion with coefficient (see also Burak and Fiete, 2012)

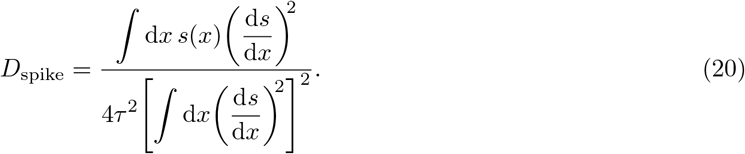

As before, *s* is the baseline firing rate configuration without noise and drive. Through the relationships in Eqs. 11 and 12, *D*_spike_ scales with *M* and *N* in the same way as *D*_input_ does:

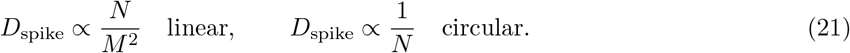

These findings are presented in Fig. 5C, D along with simulation results that confirm our theory. Spiking noise behaves similarly to input noise. Increasing bump number improves robustness to noise under linear mapping but has almost no effect under circular mapping. Bump diffusion in larger networks is exacerbated under linear mapping but suppressed under circular mapping. For both input noise and spiking noise, the conversion factor in Eq. 12 produces the differences between the two mappings. Coupling strength rescaling in Eq. 13 does not play a role because *γ* does not appear in Eqs. 10 and 20.

To evaluate noise robustness a different way, we perform mutual information analysis of networks with input noise. Mutual information describes how knowledge of one random variable can reduce the uncertainty in another, and it serves as a general metric for encoding quality. Here, we calculate the mutual information *I* between the physical coordinate encoded by the noisy network, represented by the random variable *U* with discretized sample space *U*, and the activity of a single neuron at network position *x*, represented by the random variable *S* with discretized sample space *S* (see Simulation methods):

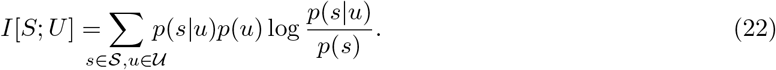

We then average *I* across neurons. Larger mean mutual information implies more precise coordinate encoding and greater robustness to noise. Thus, we expect that networks with less diffusion in Fig. 5A, B should generally contain more information. Note that the joint activities of all the neurons confer much more coordinate information than single-neuron activities do, but since estimating high-dimensional probability distributions over the former is computationally very costly, we use the latter as a metric for network performance.

The physical coordinate *U* is either position or orientation and obeys the mappings described in Fig. 4 across bump numbers *M* and network sizes *N*. To obtain the probability distributions in Eq. 22 required to calculate *I*, we initialize multiple replicate simulations at evenly spaced coordinate values *u* (Fig. 6A). We do not apply an input drive, so the networks should still encode their initialized coordinates at the end of the simulation. However, they contain input noise that degrades their encoding. Collecting the final firing rates *s*(*x*) produces *p*(*s*|*u*) for each neuron *x*. For both position and orientation, we consider narrow and wide coordinate ranges to assess network performance in both regimes.

**Figure 6:**
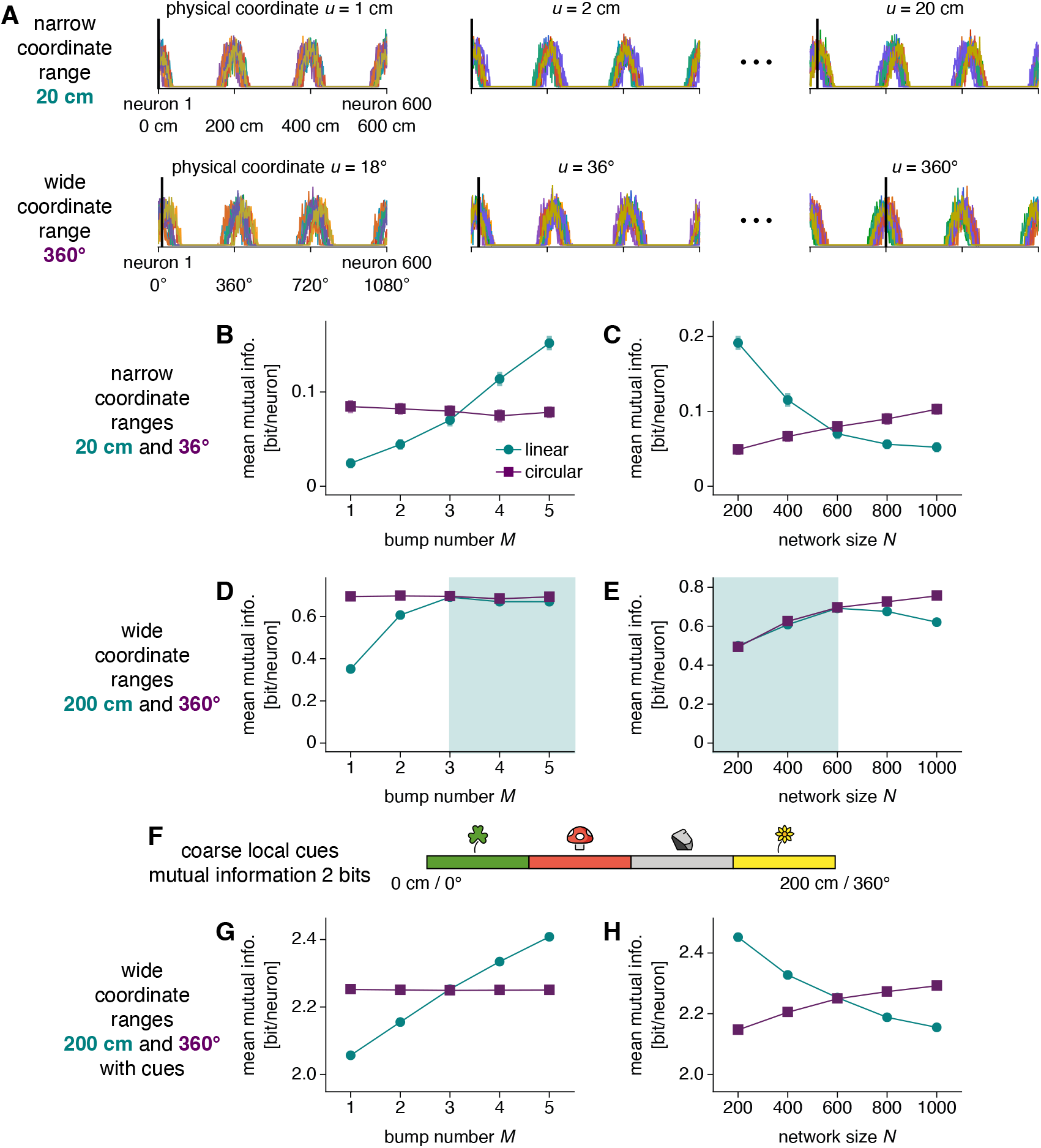
Mutual information between neural activity and physical coordinates with input noise of magnitude *σ* = 0.5. (**A**) To calculate mutual information, we initialize replicate simulations without input drive at different coordinate values (thick black lines) and record the final neural activities (thin colored lines). The physical coordinate can be linear or circular and its range can be narrow or wide; here, we illustrate two possibilities for networks with 600 neurons and 3 bumps. (**B, C**) Mutual information between physical coordinate and single-neuron activity under narrow coordinate ranges. (**B**) Information decreases with bump number for linear coordinates and remains largely constant for circular coordinates. Networks with 600 neurons. (**C**) Information decreases with network size for linear coordinates and increases for circular coordinates. Networks with 3 bumps. (**D, E**) Mutual information between physical coordinate and single-neuron activity under wide coordinate ranges. The trends in **B, C** are preserved for circular coordinates. They are also preserved for linear coordinates, except for the shaded regions in which the coordinate range exceeds the bump distance. (**F**) Coarse local cues are active over different quadrants of the wide coordinate ranges. (**G, H**) Mutual information between physical coordinate and the joint activities of a single neuron with the four cues in **F** under wide coordinate ranges. The trends in **B, C** are preserved for both linear and circular coordinates. Points indicate data from 96 replicate simulations at each coordinate value averaged over neurons and bars indicate bootstrapped standard errors of the mean.

We first consider narrow coordinate ranges. For linear coordinates, information increases as a function of *M* but decreases as a function of *N*; for circular coordinates, it does not strongly depend on *M* and increases as a function of *N* (Fig. 6B, C). These results exactly corroborate those in Fig. 5A, B obtained for bump diffusion, since we expect information and diffusion to be inversely related.

We next consider wide coordinate ranges. Our ring networks can only uniquely represent coordinate ranges up to their bump distances (converted to physical distances by Fig. 4). Beyond these values, two physical coordinates separated by the converted bump distance cannot be distinguished by the network. Our mutual information analysis captures this phenomenon; for linear coordinates, the increase in information with larger *M* or smaller *N* as observed in Fig. 6B, C disappears once the converted bump distance drops below the physical range of 200 cm (green shaded regions of Fig. 6D, E). In this regime, the benefits of more bumps and smaller networks toward decreasing diffusion are counteracted by bump ambiguity. In contrast, the circular mapping in Fig. 4 lacks bump ambiguity since the bump distance is always converted to the maximum physical range of 360°, so the same qualitative trends in mutual information are observed for any coordinate range (Fig. 6D, E).

For linear coordinates with wide ranges, the advantages of increasing bump number can be restored by coarse local cues. We illustrate this process by introducing four cues, each of which is active over a different quadrant and is otherwise inactive (Fig. 6F). They can be conceptualized as two-state sensory neurons or neural populations that fire in the presence of a local stimulus. By themselves, the cues do not encode precise coordinate values. Mutual information calculated with the joint activity of each neuron with these cues recovers the behavior observed for narrow ranges across all *M* and *N* (Fig. 6G, H). Ring attractor neurons provide information beyond the 2 bits conveyed by the cues alone, and for position, this additional information increases with more bumps and fewer neurons without saturating.

In summary, our conclusions about robustness to input noise obtained by diffusion analysis are also supported by mutual information analysis. Moreover, the latter explicitly reveals how networks encoding wide, linear coordinate ranges can leverage coarse local cues to address ambiguities and preserve the advantages of multiple bumps.

### More bumps improve robustness to connectivity noise under linear mapping

Another source of noise in biological CANs is inhomogeneity in the connectivity *W*. Perfect continuous attractor dynamics requires *W* to be invariant to translations along the network (Skaggs et al., 1995; Zhang, 1996; Samsonovich and McNaughton, 1997; Fuhs and Touretzky, 2006; Burak and Fiete, 2009), a concept related to Goldstone’s theorem in physics (Nambu, 1960; Goldstone, 1961). We consider the effect of replacing *W → W* + *V*, where *V* is a noisy connectivity matrix whose entries are independently drawn from a zeromean Gaussian distribution. *V* disrupts the symmetries of *W*. This noise is quenched and does not change over the course of the simulation, in contrast to input and spiking noise which are independently sampled in time. *V* creates a noise term

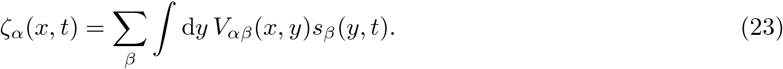

Equation 23 shows that *V* produces correlated *ζ*’s across neurons, which also differs from input and spiking noise. Because of these distinctions, the dominant effect of connectivity noise is not diffusion, but drift. *V* induces bumps to move with velocity *v*_conn_(*θ*), even without drive *b*:

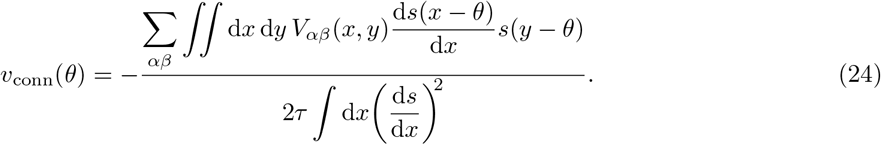

The movement is coherent but irregular, as it depends on the bump position *θ* (Fig. 7A). Itskov et al. (2011) and Seeholzer et al. (2019) refer to *v*_conn_(*θ*) as the drift velocity.

**Figure 7:**
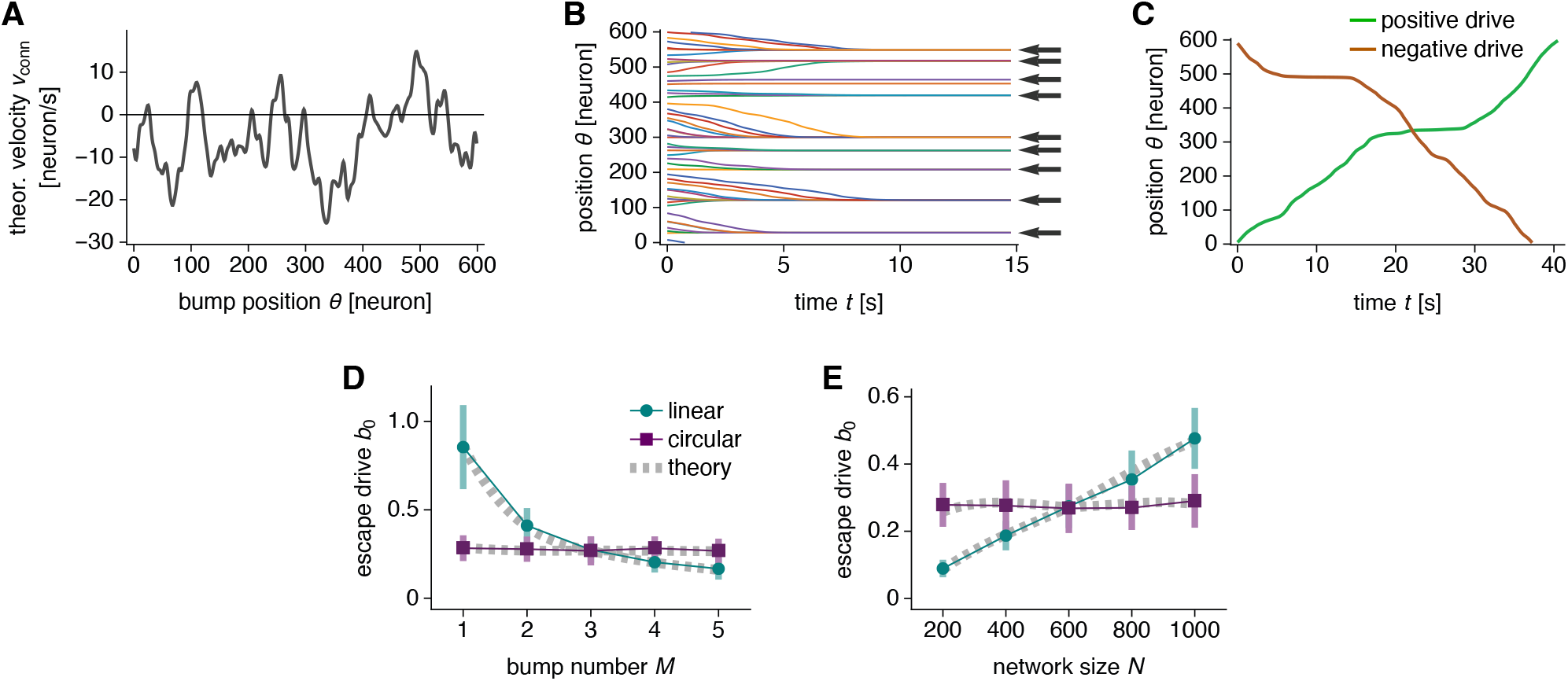
Bump trapping due to connectivity noise at low drive. (**A–C**) Networks with 600 neurons, 1 bump, and the same realization of connectivity noise of magnitude 0.002. (**A**) Theoretical values for drift velocity as a function of bump position using Eq. 24. (**B**) Bumps drift towards trapped positions over time. The drive is *b* = 0. Arrows indicate predictions from *v*_conn_(*θ*) crossing 0 with negative slope in **A**. Lines indicate simulations with different starting positions. (**C**) Bump trajectories with smallest positive and negative drive required to travel through the entire network. Respectively, *b* = 0.75 and *b* = *−* 0.52. The larger of the two in magnitude is the escape drive *b*_0_ = 0.75. Note that positions with low bump speed exhibit large velocities in the opposite direction in **A**. (**D, E**) Networks with multiple realizations of connectivity noise of magnitude 0.002. (**D**) Escape drive decreases with bump number under linear mapping and remains largely constant under circular mapping. Networks with 600 neurons. (**E**) Escape drive increases with network size under linear mapping and remains largely constant under circular mapping. Networks with 3 bumps. Points indicate simulation means over 48 realizations and bars indicate standard deviations. Dotted gray lines indicate Eq. 26 averaged over 96 realizations.

Connectivity noise traps bumps at low drive *b*. We first consider *b* = 0, so bump motion is governed solely by drift according to d*θ/*d*t* = *v*_conn_(*θ*). The bump position *θ* has stable fixed points wherever *v*_conn_(*θ*) crosses 0 with negative slope (Itskov et al., 2011; Seeholzer et al., 2019). Simulations confirm that bumps drift towards these points (Fig. 7B). The introduction of *b* adds a constant *v*_drive_ that moves the curve in Fig. 7A up for positive *b* or down for negative *b*:

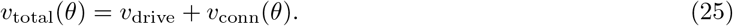

If *v*_total_(*θ*) still crosses 0, bumps would still be trapped. The absence of bump motion in response to coordinate changes encoded by *b* would be a catastrophic failure of path integration. To permit bump motion through the entire network, the drive must be strong enough to eliminate all zero-crossings. Figure 7C shows bump motion at this drive for both directions of motion. The positive *b* is just large enough for the bump to pass through the region with the most negative *v*_conn_(*θ*) in Fig. 7A; likewise for negative *b* and positive *v*_conn_(*θ*). We call the larger absolute value of these two drives the escape drive *b*_0_. Simulations show that *b*_0_ decreases with bump number *M* and increases with network size *N* under linear mapping (Fig. 7D, E). A smaller *b*_0_ implies weaker trapping, so smaller networks with more bumps are more resilient against this phenomenon. Under circular mapping, however, *b*_0_ demonstrates no significant dependence on *M* or *N*. We can predict *b*_0_ by inverting the relationship in Eq. 8 between *b* and *v*:

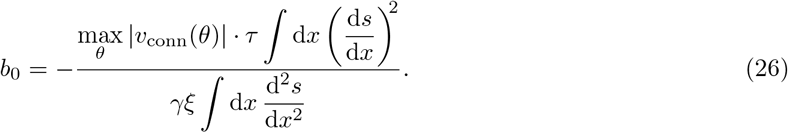

This theoretical result agrees well with values obtained by simulation (Fig. 7D, E). In the Theoretical model section, we present a heuristic argument (Eq. 124) that leads to the observed scaling of escape drive on *M* and *N* :

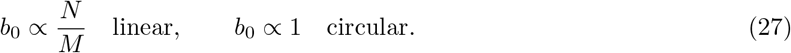

At high drive |*b*| *> b*_0_, attractor bumps are no longer trapped by the drift velocity *v*_conn_(*θ*). Instead, the drift term produces irregularities in the total velocity *v*_total_(*θ*) (Fig. 8A). They can be decomposed into two components: irregularities between directions of motion and irregularities within each direction. Both imply errors in path integration because *v* and *b* are not strictly proportional. To quantify these components, we call |*v*_+_(*θ*)| and |*v*_*−*_(*θ*)| the observed bump speeds under positive and negative *b*. We define speed difference as the unsigned difference between mean speeds in either direction, normalized by the overall mean speed:

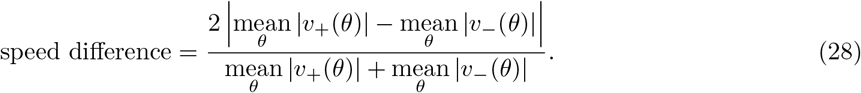

We then define speed variability as the standard deviation of speeds within each direction, averaged over both directions and normalized by the overall mean speed:

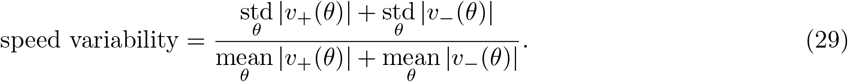

Speed difference and speed variability follow the same trends under changes in bump number *M* and network size *N* (Fig. 8B–E). Under linear mapping, they decrease with *M* and increase with *N*. Under circular mapping, they do not significantly depend on *M* and *N*. These are also the same trends exhibited by the escape drive *b*_0_ (Fig. 7D, E). In terms of theoretical quantities, the formulas for speed difference and variability become

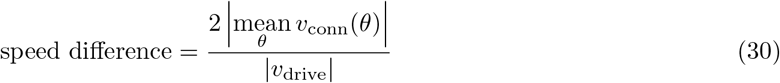

and

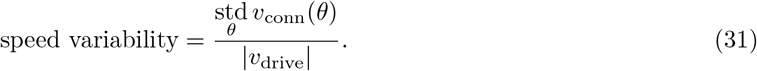

These expressions match the observed values well (Fig. 8B–E). In the Theoretical methods section, we calculate the observed dependence of speed difference (Eq. 113) and variability (Eq. 120) on *M* and *N*:

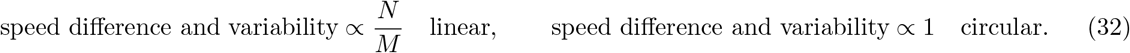

**Figure 8:**
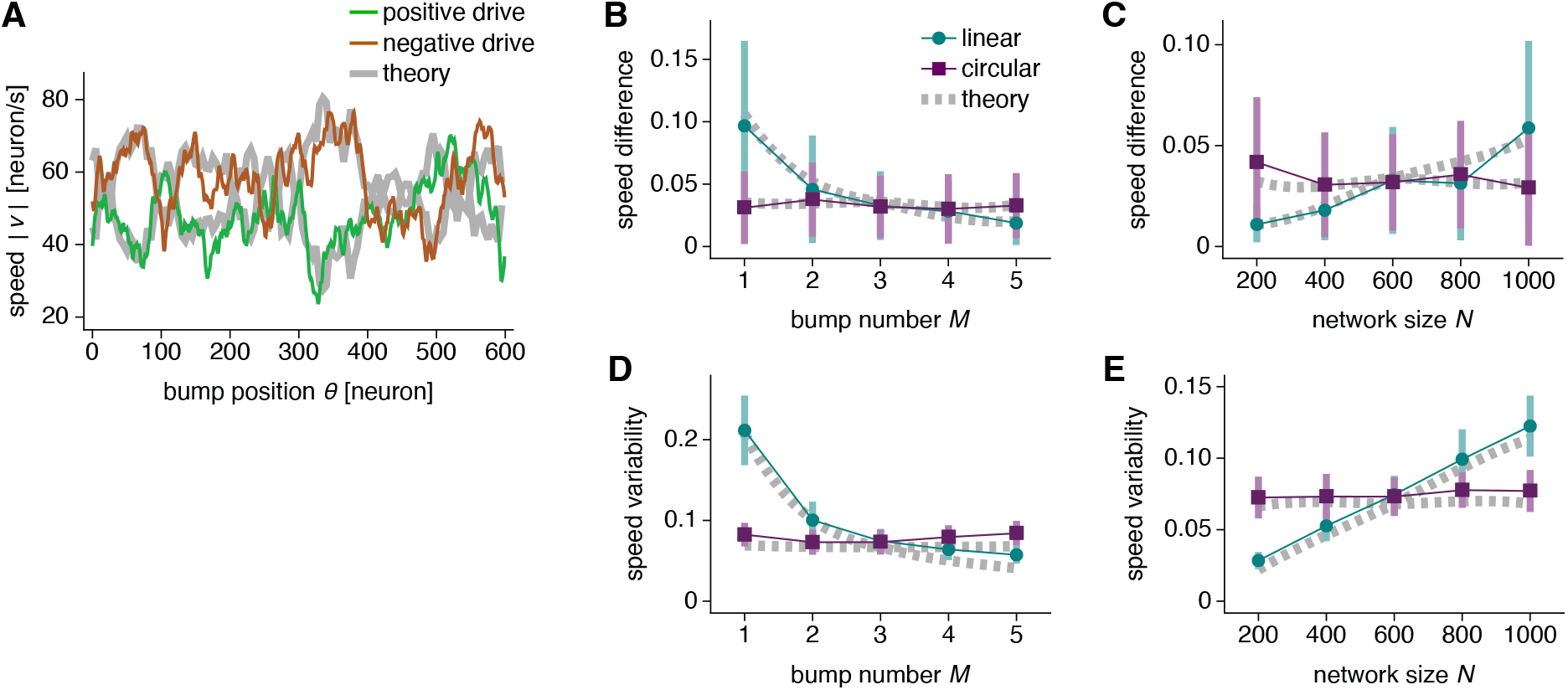
Bump speed irregularity due to connectivity noise at high drive. (**A**) Bump speed as a function of bump position with connectivity noise of magnitude 0.002 and drive *b* = 1.5. Network with 600 neurons, 1 bump, and the same realization of connectivity noise as in Fig. 7A–C. Thick gray lines indicate Eq. 25. (**B–E**) Networks with multiple realizations of connectivity noise of magnitude 0.002 and drive *b* = 1.5. (**B**) Speed difference between directions decreases with bump number under linear mapping and remains largely constant under circular mapping. Networks with 600 neurons. (**C**) Speed difference increases with network size under linear mapping and remains largely constant under circular mapping. Networks with 3 bumps. (**D, E**) Same as **B, C**, but for speed variability within each direction. Points indicate simulation means over 48 realizations and bars indicate standard deviations. Dotted gray lines indicate Eqs. 30 and 31 averaged over 96 realizations.

For all results related to connectivity noise, the coupling strength rescaling in Eq. 13 produces the differences between the two mappings via the *γ* in Eq. 8. The conversion factor in Eq. 12 does not play a role because escape drive, speed difference, and speed variability do not have dimensions of network distance.

To summarize, CANs with imperfect connectivity benefit from more attractor bumps when encoding linear coordinates. This advantage is present at all driving inputs and may be more crucial for larger networks. On the other hand, connectivity noise has largely the same consequences for networks of all bump numbers and sizes when encoding circular coordinates.

## Discussion

We demonstrated how CANs capable of path integration respond to three types of noise. Additive synaptic input noise and Poisson spiking noise cause bumps to diffuse away from the coherent motion responsible for path integration (Figs. 3 and 5). This diffusion is accompanied by a decrease in mutual information between neural activity and encoded coordinate (Fig. 6). Connectivity noise produces a drift velocity field that also impairs path integration by trapping bumps at low drive and perturbing bump motion at high drive (Figs. 7 and 8).

For all three types of noise, CANs with more attractor bumps exhibit less deviation in bump motion in network units. This is observed across network parameters (Figs. 9 and 10 in the Appendix). Thus, CANs can more robustly encode linear variables whose mapping inherits network units and does not rescale with bump number (Fig. 4A). If grid cell networks were to encode spatial position in this manner, then multiple attractor bumps would be preferred over a single bump. Gu et al. (2018) report experimental evidence supporting multi-bump grid networks obtained by calcium imaging of mouse medial entorhinal cortex. Our work implies that the evolution of such networks may have been partially encouraged by biological noise. Additional bumps introduce greater ambiguity among positions encoded by each bump, but this can be resolved by a rough estimate of position from additional cues, such as local landmarks (O’Keefe and Burgess, 2005; Krupic et al., 2014; Bush and Burgess, 2014; Hardcastle et al., 2015), another grid module with different periodicity (O’Keefe and Burgess, 2005; McNaughton et al., 2006; Stensola et al., 2012; Stemmler et al., 2015; Kang and Balasubramanian, 2019; Khona et al., 2022), or a Bayesian prior based on recent memory (Sreenivasan and Fiete, 2011). In this way, grid modules with finer resolution and more attractor bumps could maintain a precise egocentric encoding of position, while coarser modules and occasional allocentric cues would identify the true position out of the few possibilities represented. We explicitly explored one realization of this concept and observed how cues enable networks to improve their information content by increasing bump number, despite a concomitant increase in bump ambiguity (Fig. 6F–H).

**Figure 9:**
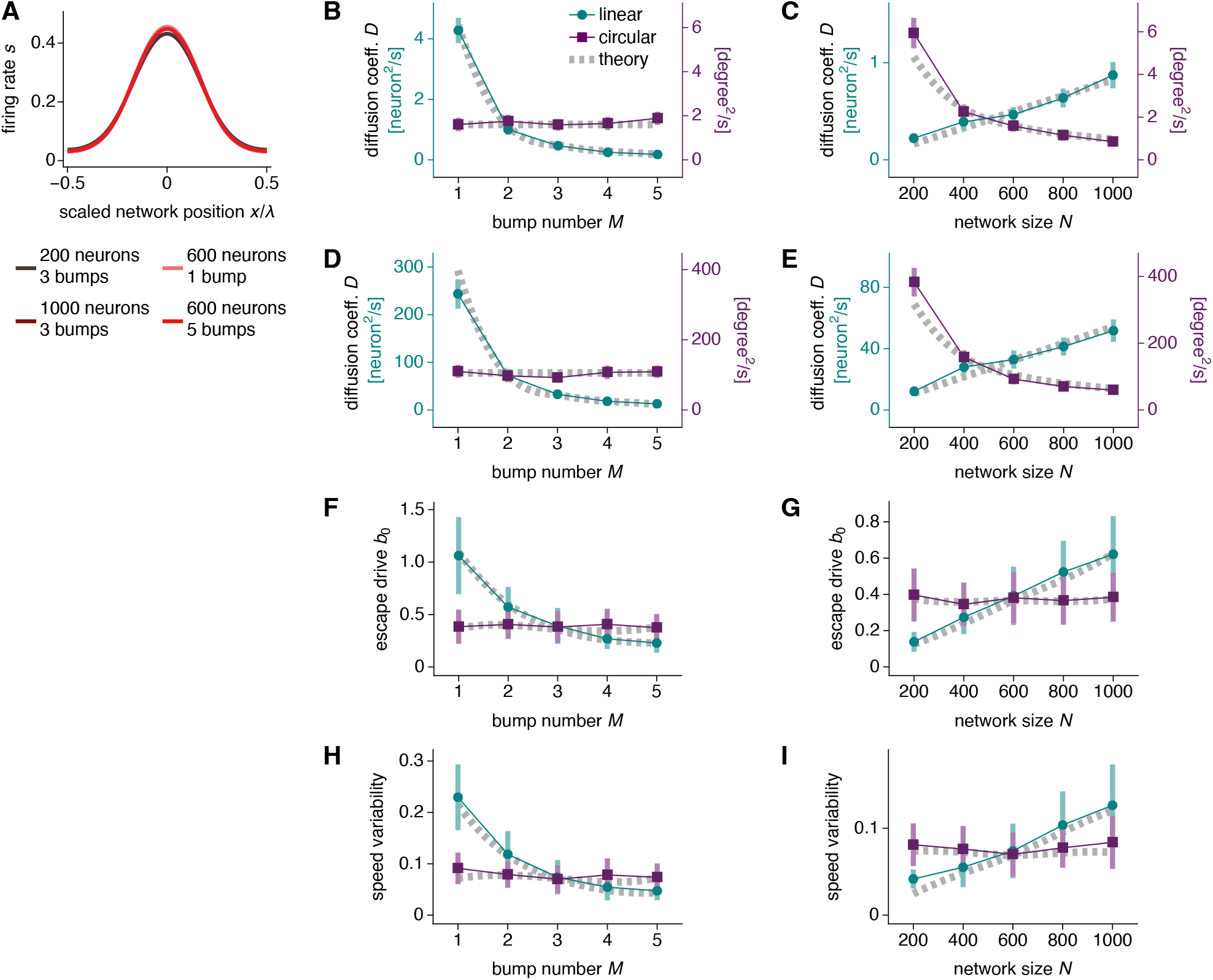
Main results repeated for networks with a logistic activation function. (**A**) The scaled bump shape remains invariant across network sizes and bump numbers, accomplished by rescaling connectivity strengths according to Eq. 7. Curves for different parameters lie over one another. (**B, C**) Networks with synaptic input noise. Bump diffusion follows the same qualitative behavior as in Fig. 5A, B. (**D, E**) Networks with Poisson spiking noise. Bump diffusion follows the same qualitative behavior as in Fig. 5C, D. (**F–I**) Networks with connectivity noise. (**F, G**) Escape drive follows the same qualitative behavior as in Fig. 7D, E. (**F, G**) Bump speed variability follows the same qualitative behavior as in Fig. 8D, E. The activation function *ϕ* takes the form in Eq. 134. In **F**–**I**, we use connectivity noise of magnitude 0.003. In **H, I**, we use drive *b* = 2.5. The rest of the parameters are identical in value to those used in the main text.

**Figure 10:**
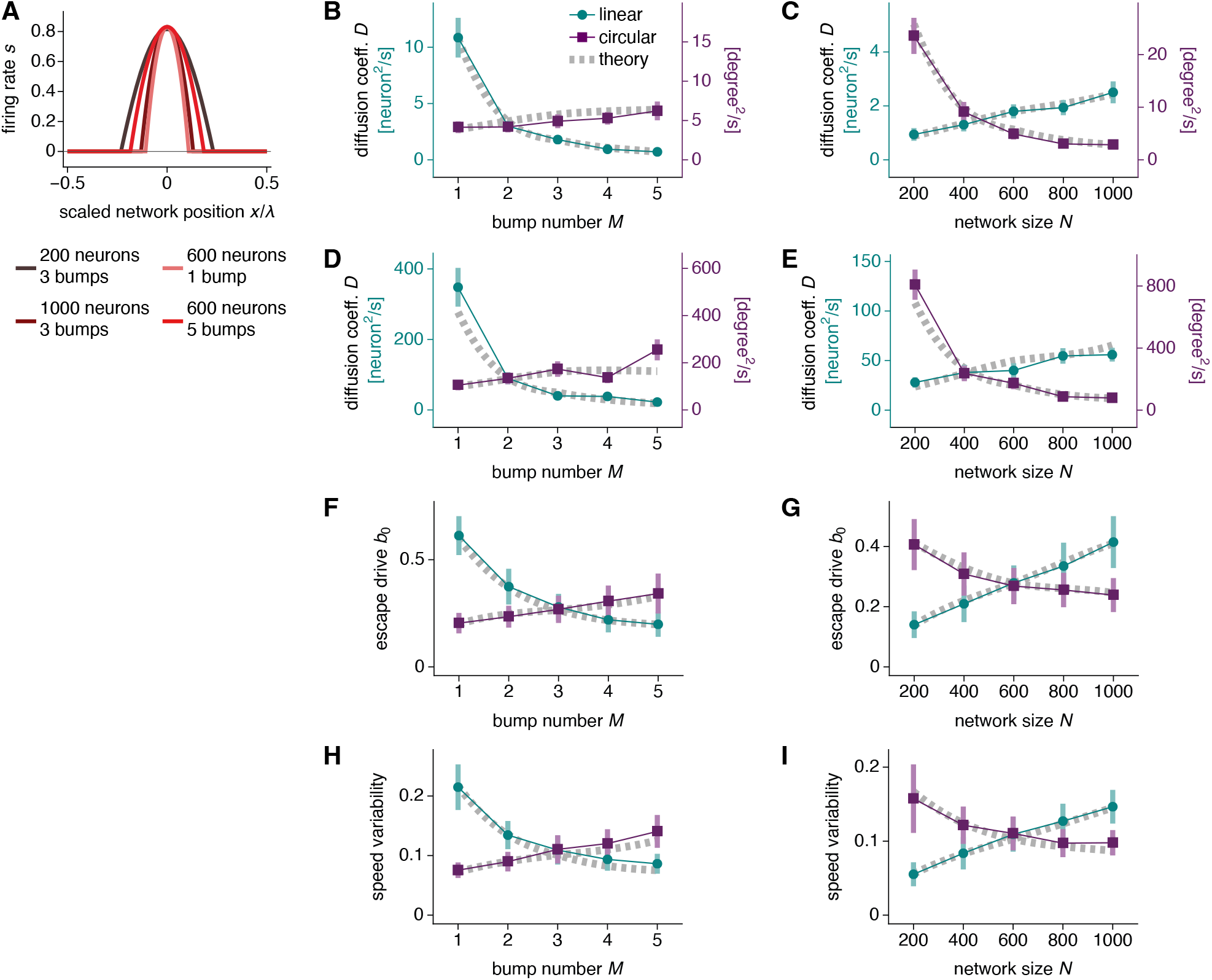
Main results repeated for networks without rescaling of connectivity strengths according to Eq. 7. (**A**) The scaled bump shape no longer remains invariant across network sizes and bump numbers. (**B, C**) Networks with synaptic input noise. Bump diffusion follows the same qualitative behavior as in Fig. 5A, B, except that here it slightly increases with bump number under circular mapping. (**D, E**) Networks with Poisson spiking noise. Bump diffusion follows the same qualitative behavior as in Fig. 5C, D, except that here it slightly increases with bump number under circular mapping. (**F–I**) Networks with connectivity noise. (**F, G**) Escape drive follows the same qualitative behavior as in Fig. 7D, E under linear mapping. It slightly increases with bump number and decreases with network size under circular mapping. (**H, I**) Bump speed variability follows the same qualitative behavior as in Fig. 8D, E under linear mapping. It slightly increases with bump number and decreases with network size under circular mapping. The connectivity *W* still takes the form in Eq. 38, except that here we fix *w* = 0.04 across all bump numbers and network sizes. In **H, I**, we use drive *b* = 1.0. The rest of the parameters are identical in value to those used in the main text.

In contrast, CANs encoding circular variables may rescale under different bump numbers to match periodicities (Fig. 4B), which eliminates any influence of bump number on encoding accuracy for all three types of noise. If head direction networks were to encode orientation in this manner, then they would face less selective pressure to evolve beyond the single-bump configuration observed in *Drosophila* (Seelig and Jayaraman, 2015). Moreover, without the assumption of bump shape invariance accomplished by Eq. 7, robustness to all three types of noise decreases with bump number, which actively favors single-bump orientation networks (Fig. 10 in the Appendix). Further experimental characterization of bump number in biological CANs, perhaps through techniques proposed by Widloski et al. (2018), would test the degree to which the brain can leverage the theoretical advantages identified in this work.

Under linear mapping, larger CANs exhibit more errors in path integration from all three types of noise. The immediate biological implication is that larger brains face a dramatic degradation of CAN performance, accentuating the importance of suppressing error with multi-bump networks. However, this simple rule that one neuron always represents a fixed physical interval does not need to be followed. Furthermore, larger animals may tolerate greater absolute errors in path integration because they interact with their environments over larger scales. Nevertheless, our results highlight the importance of considering network size when studying the performance of noisy CANs. Under circular mapping, bump diffusion decreases with network size for input and spiking noise, and the magnitude of errors due to connectivity noise is independent of network size. This implies that head direction networks can benefit from incorporating more neurons; the observed interactions between such networks across different mammalian brain regions may act in this manner to suppress noise (Taube, 2007).

The computational advantages of periodic over nonperiodic encodings has been extensively studied in the context of grid cells (Fiete et al., 2008; Sreenivasan and Fiete, 2011; Mathis et al., 2012; Wei et al., 2012; Almeida et al., 2015; Wei et al., 2015; Stemmler et al., 2015). Our results extend these findings by demonstrating that some kinds of periodic encodings can perform better than others. Our results also contribute to a rich literature on noisy CANs. Previous studies have investigated additive input noise (Compte et al., 2000; Wu et al., 2008; Burak and Fiete, 2012; Kilpatrick and Ermentrout, 2013; Seeholzer et al., 2019), multiplicative input noise (Krishnan et al., 2018), spiking noise (Burak and Fiete, 2009, 2012; Wei et al., 2012; Almeida et al., 2015; Bouchacourt and Buschman, 2019; Seeholzer et al., 2019), and quenched noise due to connectivity or input inhomogeneities (Zhang, 1996; Itskov et al., 2011; Kilpatrick and Ermentrout, 2013; Seeholzer et al., 2019; Can and Krishnamurthy, 2021). Among these works, the relationship between bump number and noise has only been considered in the context of multiple-item working memory, in which each network can be loaded with various numbers of bumps (Wei et al., 2012; Almeida et al., 2015; Krishnan et al., 2018; Bouchacourt and Buschman, 2019). Interestingly, they find that robustness to noise decreases with bump number, which is opposite to our results (cf. Almeida et al., 2015, who report no dependence between bump number and encoding accuracy under certain conditions). It appears that CANs designed for path integration with fixed bump number and CANs designed for multiple-item working memory with variable bump number differ crucially in their responses to noise. Further lines of investigation that compare these two classes would greatly contribute to our general understanding of CANs.

Beyond our concrete results on CAN performance, our work offers a comprehensive theoretical framework for studying path-integrating CANs. We derive a formula for the multi-bump attractor state and a Lyapunov functional that governs its formation. We calculate all key dynamical quantities such as velocities and diffusion coefficients in terms of firing rates. Our formulas yield scaling relationships that facilitate an intuitive understanding for their dependence on bump number and network size. Much of our theoretical development does not assume a specific connectivity matrix or nonlinear activation function, which allows our results to have wide significance. For example, we expect them to hold for path-integrating networks that contain excitatory synapses. Other theories have been developed for bump shape (Wu et al., 2002; Xie et al., 2002; Itskov et al., 2011; Kilpatrick and Ermentrout, 2013; Widloski, 2015; Krishnan et al., 2018), path integration velocity (Xie et al., 2002; Mosheiff and Burak, 2019), diffusion coefficients (Wu et al., 2008; Burak and Fiete, 2012; Kilpatrick and Ermentrout, 2013; Krishnan et al., 2018; Seeholzer et al., 2019), and drift velocity (Zhang, 1996; Itskov et al., 2011; Seeholzer et al., 2019). Our work unifies these studies through a simple framework that features path integration, multiple bumps, and a noise term that can represent a wide range of sources. It can be easily extended to include other components of theoretical or biological significance, such as slowly-varying inputs (Tsodyks and Sejnowski, 1995; Fung et al., 2010; Kilpatrick and Ermentrout, 2013), synaptic plasticity (Stringer et al., 2002; Renart et al., 2003), neural oscillations (Thurley et al., 2008; Navratilova et al., 2012; Kang and DeWeese, 2019), and higher-dimensional attractor manifolds (Ermentrout and Cowan, 1979; Samsonovich and McNaughton, 1997).

## Theoretical model

### Architecture

We investigate CAN dynamics through a 1D ring attractor network. This class of network has been analyzed in previous theoretical works, and at various points, our calculations will parallel those in Xie et al. (2002); Itskov et al. (2011); Burak and Fiete (2012); Kilpatrick and Ermentrout (2013); Widloski (2015); Seeholzer et al. (2019); Mosheiff and Burak (2019).

There are two neurons at each position *i* = 0, …, *N* − 1 with population indexed by *α* ∈ {L, R} (Fig. 1A). For convenient calculation, we unwrap the ring and connect copies end-to-end, forming a linear network with continuous positions *x* ∈ (*−∞,∞*). Unless otherwise specified, integrals are performed over the entire range. To map back onto the finite-sized ring network, we enforce our results to have a periodicity *λ* that divides *N*. For example, *λ* = *N* corresponds to a single-bump configuration. Integrals would then be performed over [0, *N*), with positions outside this range corresponding to their equivalents within this range.

The network obeys the following dynamics for synaptic inputs *g*:

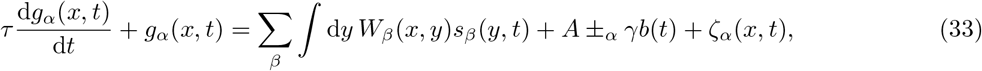

where *±*_L_ means*−* and *±*_R_ means +, and the opposite for ∓_*α*_. *τ* is the neural time constant, *W* is the synaptic connectivity, and *A* is the resting input. The nonlinear activation function *ϕ* converts synaptic inputs to firing rates:

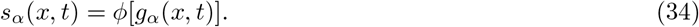

Most of our results will apply to general *ϕ*, but we also consider a ReLU activation function specifically:

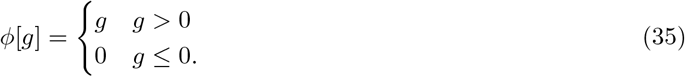

In this section, we will explicitly mention when we specifically consider the ReLU case, and we will always simplify the function away. Thus, if an expression contains the symbol *ϕ*, then it applies to general *ϕ*. In the Results section, formulas and scalings for *D*_input_, *D*_spike_, *v*_drive_, and *v*_conn_(*θ*), as well as all simulation results invoke Eq. 35. We will use this form in the Bump shape *g* subsection *b* is the driving input, *γ* is its coupling strength, and *ζ* is the noise, which can take different forms. *γb* and *ζ* are small compared to the rest of the right-hand side of Eq. 33. For notational convenience, we will often suppress dependence on *t*.

*W*_*β*_(*x, y*) obeys a standard continuous attractor architecture based on a symmetric and translation invariant *W* :

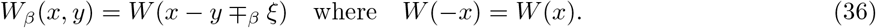

Each population *β* deviates from *W* by a small shift *ξ* ≪ *N* in synaptic outputs. Thus, the following approximation holds:

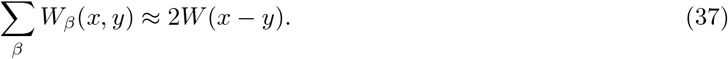

We will consider the specific form of *W* (Fig. 1B):

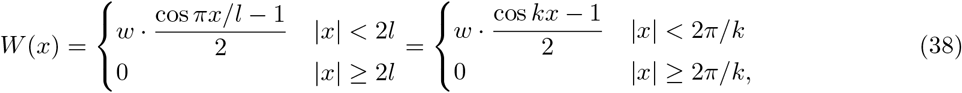

where *k* = *π/l*. We will explicitly mention when we specifically consider this form; in fact, we only do so for Eqs. 46, 47, 59, and 60, as well as for our simulation results in the Results section. Otherwise, each expression applies to general *W*.

### Baseline configuration without drive and noise

#### Linearized dynamics and bump distance *λ*

First, we consider the case of no drive *b* = 0 and no noise *ζ* = 0. The dynamical equation Eq. 33 becomes

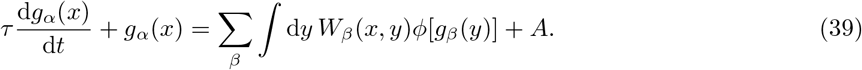

Since the right-hand side no longer depends on *α, g* must be the same for both populations, and we can use Eq. 37 to obtain

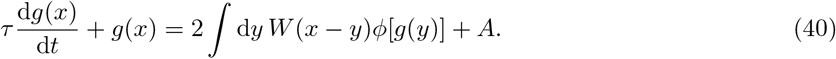

We analyze these dynamics using the Fourier transform *F*. Our chosen convention, applied to the function *h*, is

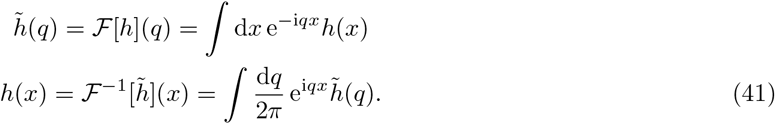

Fourier modes 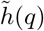 represent sinusoids with wavenumber *q* and corresponding wavelength 2*π/q*. Applying this transform to Eq. 40, we obtain

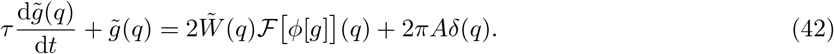

In this subsection, we consider the case of small deviations, such that *g*(*x*) *≈ g*_0_ and *ϕ*[*g*(*x*)] *≈ φ*[*g*_0_] + *ϕ*′[*g*_0_](*g*(*x*) *− g*_0_). Then, Eq. 42 becomes

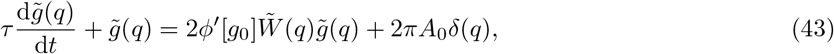

where 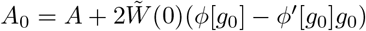. The solution to this linearized equation for *q* ≠ 0 is

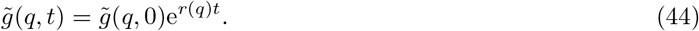

Each mode grows exponentially with rate

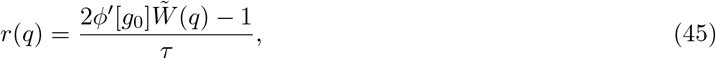

so the fastest-growing component of *g* is the one that maximizes 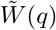, as stated in Eq. 5 of the Results section. The wavelength 2*π/q* of that component predicts the bump distance *λ*.

For the specific *W* in Eq. 38, its Fourier transform is

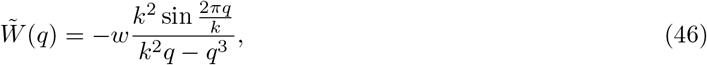

so

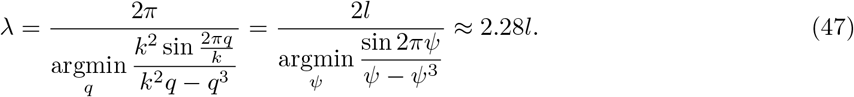

*λ* is proportional to *l*, as also noted by Fuhs and Touretzky (2006); Burak and Fiete (2009); Kang and Balasubramanian (2019); Khona et al. (2022).

#### Bump shape *g*

We call the steady-state synaptic inputs *g* without drive and noise the baseline configuration. To calculate its shape, we must account for the nonlinearity of the activation function *ϕ* and return to Eq. 42. We invoke our particular form of *ϕ* in Eq. 35 to calculate ℱ[*ϕ*[*g*]] (*q*). *g* must be periodic, and its periodicity is the bump distance *λ* with wavenumber *κ* = 2*π/λ*. Without loss of generality, we take *g* to have a bump centered at 0. Since *W* is symmetric, *g* is an even function. We define *z* as the position where *g* crosses 0:

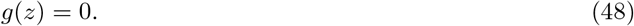

If *g* is approximately sinusoidal, then *g*(*x*) *>* 0 wherever *nλ − z < x < nλ* + *z* for any integer *n*. The ReLU activation function in Eq. 35 implies

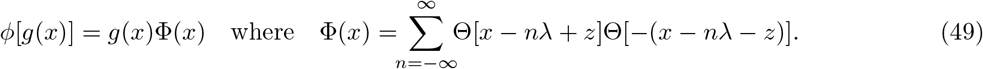

Θ is the Heaviside step function. The Fourier transform for Φ is

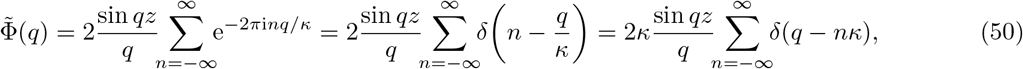

where the second equality comes from the Fourier series for a Dirac comb. Therefore,

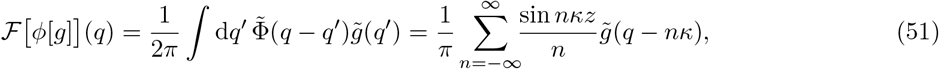

so Eq. 42 becomes

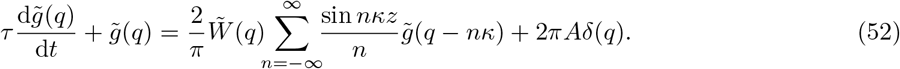

This equation describes the full dynamics of *g* with a ReLU activation function. It contains couplings between all modes *q* that are multiples of the wavenumber *κ*, which corresponds to the bump distance.

To find the baseline *g*, we set d 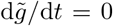 We also simplify 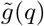 by only considering the lowest modes that couple to each other: *q* = 0, *±κ*. Due to symmetry, 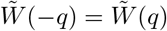 and 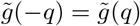. Eq. 52 gives

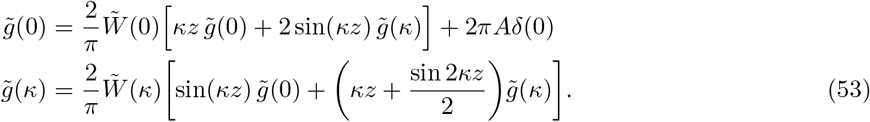

Now we need to impose Eq. 48: *g*(*z*) = 0. To do so, we note that 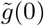 and 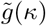 are both proportional to *δ*(0) according to Eq. 53. That means 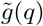 has the form

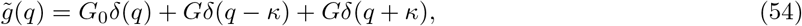

where *G*_0_ and *G* are the Fourier modes with delta functions separated. This implies

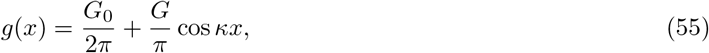

and *g*(*z*) = 0 implies

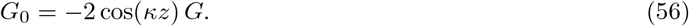

Substituting Eqs. 54 and 56 into Eq. 53, we obtain

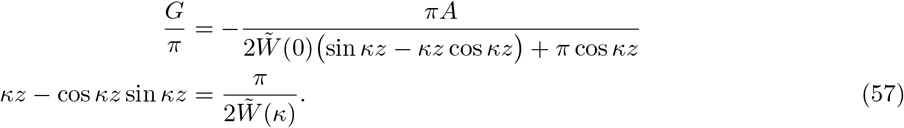

We can solve the second equation of Eq. 57 for *κz* and then substitute it into the first equation to obtain *G*. This then gives us *g*(*x*), which becomes through Eqs. 55 and 56

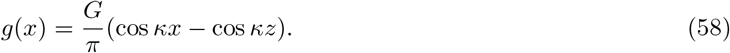

In particular, let’s use the *W* defined by Eq. 38 with Fourier transform Eq. 46. Then,

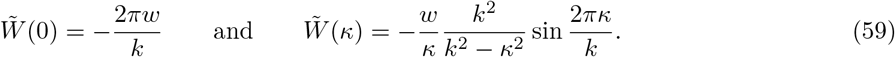

Thus,

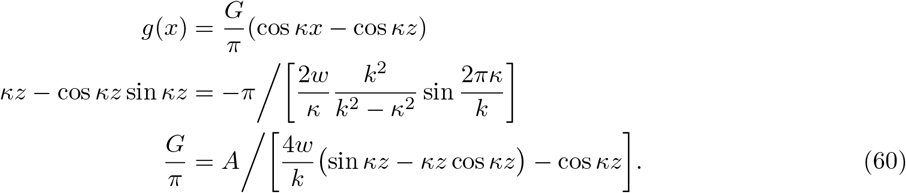

This provides expressions for *a* and *d* in Eq. 4 of the Results section, where *a* = *G/π* and *d* = *−*(*G/π*) cos *κz*.

#### Lyapunov functional and bump distance *λ*

The dynamical equation in Eq. 40 admits a Lyapunov functional. In analogy to the continuous Hopfield model (Hopfield, 1984), we can define a Lyapunov functional in terms of *s*(*x*) = *ϕ*[*g*(*x*)]:

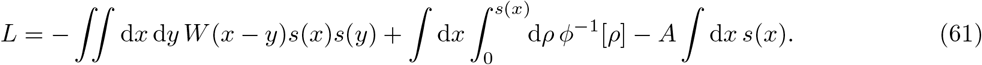

The nonlinearity *ϕ* must be invertible in the range (0, *s*) for any possible firing rate *s*. For *L* to be bounded from below for a network of any size *N*, we need

1. *W* (*x*) to be negative definite, and
2. 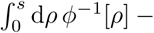 *As* to be bounded from below for any possible firing rate *s*.

We can check that these hold for our particular functions. Equation 38 immediately shows that the first condition is met. Equation 35 states that *ϕ*^*−*1^[*ρ*] = *ρ* when *ρ >* 0, so 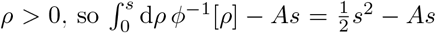, which satisfies the second condition.

Now we take the time derivative and use Eq. 40:

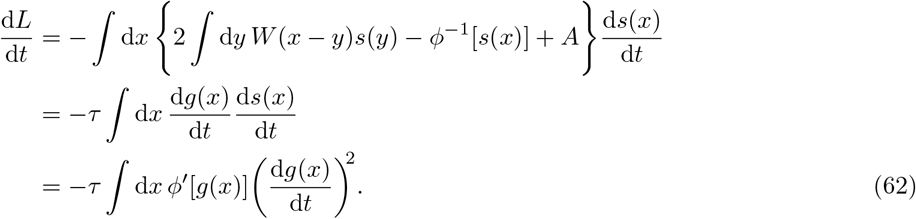

As long as *ϕ* is a monotonically nondecreasing function, d*L/*d*t* ≤ 0. Thus, *L* is a Lyapunov functional.

Now we seek to simplify Eq. 61. Suppose we are very close to a steady-state solution, so d*g/*d*t* ≈ 0. We substitute Eq. 40 into Eq. 61 to obtain

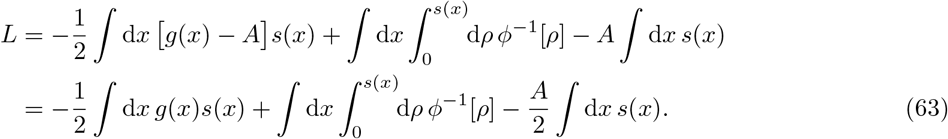

Now we invoke our ReLU *ϕ* from Eq. 35 to obtain

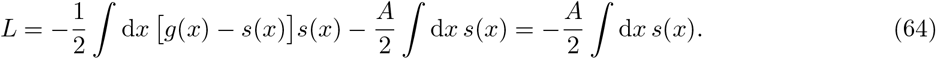

The last equality was obtained by noticing that for any *x*, either *s*(*x*) = 0 or *g*(*x*) − *s*(*x*) = 0 with our *ϕ*. Therefore, the stable solution that minimizes *L* is the one that maximizes the total firing rate.

We can apply our sinusoidal *g* in Eq. 58 to perform the integral:

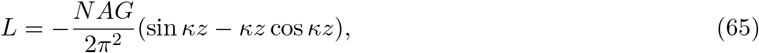

where *N* is the network size. So *L* depends on *G* and the quantity *κz*, which we will rewrite as *ψ*. We now simplify Eq. 65 using Eq. 57:

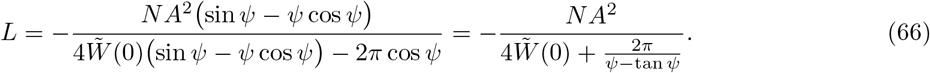

Note that 1*/*(tan *ψ − ψ*) is a monotonically increasing function of *ψ* in the range [0, *π*], so to minimize *L*, we need to minimize *ψ*. Meanwhile, Eq. 57 now reads 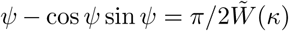. The left-hand side is also a monotonically increasing function of *ψ* in the range [0, *π*], so to minimize *ψ*, we need to maximize 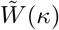. Thus, the Lyapunov stable wavelength *λ* = 2*π/κ* is the one that maximizes 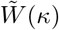. This is the same mode that grows the fastest for the linearized dynamics in Eq. 45.

### Bump motion under drive and noise

#### Dynamics along the attractor manifold

Now that we have determined the baseline configuration *g*, including the bump shape and bump distance, we investigate its motion under drive *b* and noise *ζ*. We introduce *θ* to label the position of the configuration. It can be defined as the center of mass or the point of maximum activity of one of the bumps. We expand the full time-dependent configuration with respect to the baseline configuration located at *θ*:

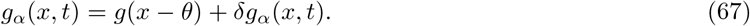

*g*(*x − θ*) solves Eq. 40 with d*g/*d*t* = 0; to facilitate calculations below, we will write the baseline equation in this form:

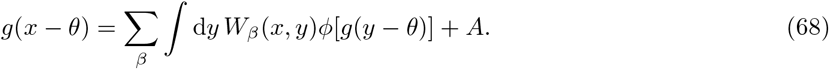

Substituting Eq. 67 into Eq. 33 and invoking Eq. 68, we obtain the following linearized dynamics for *δg*:

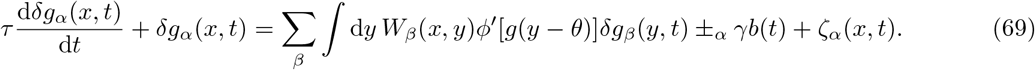

We can rewrite this as

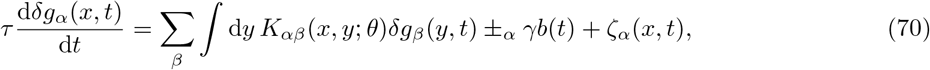

where

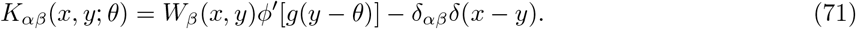

We will often suppress the argument of derivatives of *g*. If we consider a configuration located at *θ*, d*g/*d*x* implies d*g*(*x − θ*)*/*d*x*. We make the argument explicit when necessary.

If we differentiate Eq. 68 by *θ*, we obtain

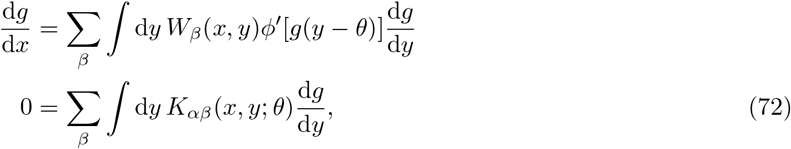

which indicates that d*g/*d*x* is a right eigenvector of *K* with eigenvalue 0. To be explicit about this, we recover the discrete case by converting continuous functions to vectors and matrices:

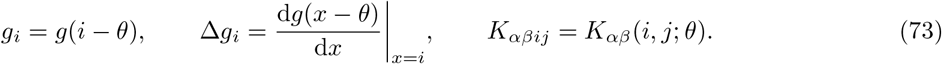

If we concatenate matrices and vectors across populations as

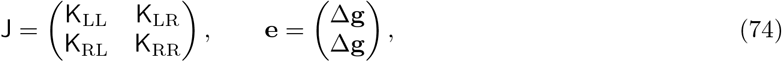

**e** is the right null eigenvector J: 0 = Σ_*j*_ *J*_*ij*_ *e*_*j*_

Since *K* is not symmetric, its left and right eigenvectors may be different. To find the left null eigenvector, we again differentiate Eq. 68 with respect to *θ*, but this time interchanging variables *x* and *y*:

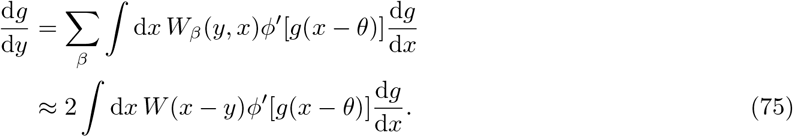

The second equality is obtained from Eqs. 36 and 37. Replacing the position *y* by *y* ± _*β*_ *ξ*, where *ξ* is the connectivity shift, we get

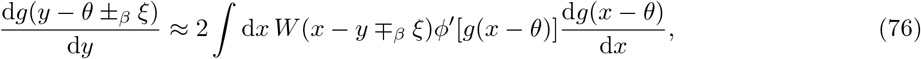

where we have made the arguments of *g* explicit. Let’s define shifted versions of the baseline *g* for each population *α*:

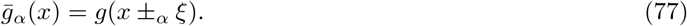

Since *ξ* is small,

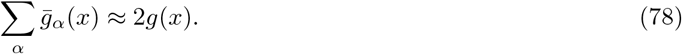

Applying these expressions to Eq. 76 and recalling Eq. 36,

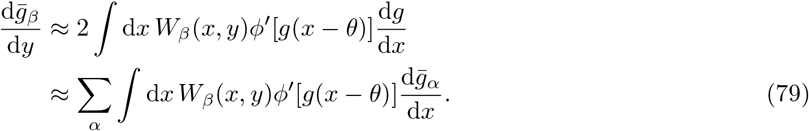

Finally, we multiply both sides of the equation by *ϕ*′[*g*(*y − θ*)] to obtain

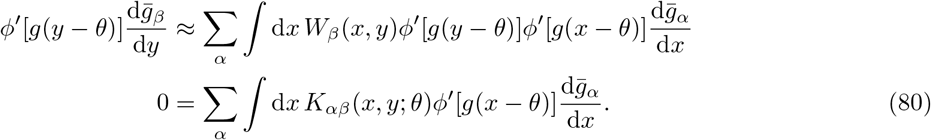

Thus 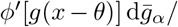 d*x* is the left null eigenvector for *K*_*αβ*_. Again, to be explicit, the discrete equivalent is

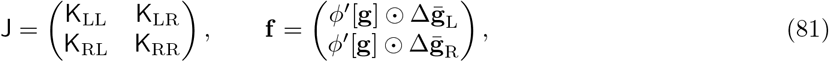

where ⊙ represents element-wise (Hadamard) multiplication. Then, **f** is the left null eigenvector of J: 0 = Σ_*i*_ *J*_*ij*_ *f*_*i*_.

We now revisit Eq. 67 and assume that *g* changes such that the bumps slowly move along the attractor manifold:

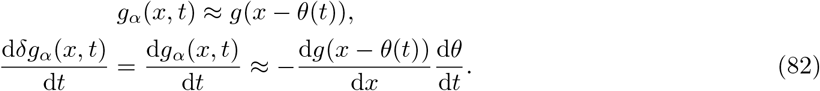

Again for simplicity, we will often suppress arguments of derivatives of *g* and dependence on *t*. We return to Eq. 70, project it along the left null eigenvector, and apply Eq. 82 to obtain

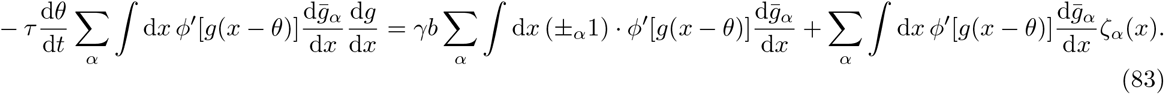

The velocity of bump motion is given by d*θ/*d*t*. It is

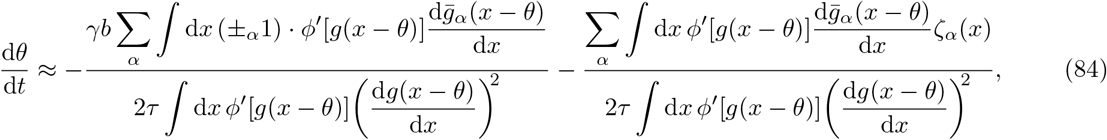

where we have made the arguments of *g* explicit. This equation encapsulates all aspects of bump motion for our theoretical model. It includes dependence on both drive *b* and noise *ζ*, the latter of which is kept in a general form. We will proceed by considering specific cases of this equation.

#### Path integration velocity *v*_drive_ due to driving input *b*

The noiseless case of Eq. 84 with *ζ*_*α*_(*x*) = 0 yields the bump velocity due to drive *b*, which is responsible for path integration:

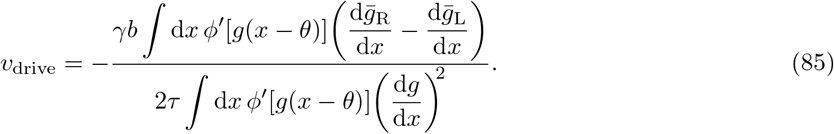

Note that this expression is independent of the position *θ*. We can explicitly remove *θ* by shifting the dummy variable *x → x* + *θ*:

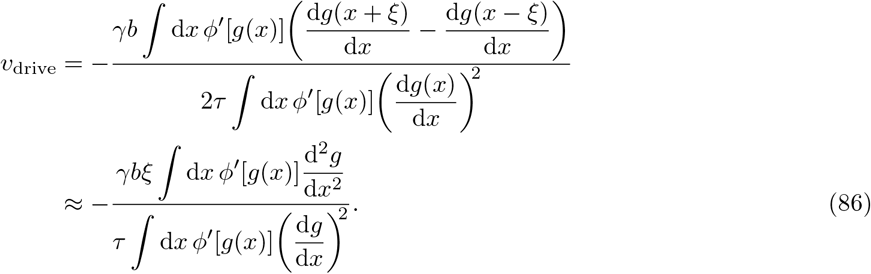

Now let’s consider the specific ReLU activation function *ϕ*. Equation 35 implies

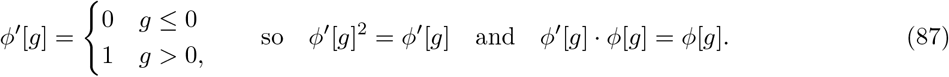

These identities, along with the definition for *s* (Eq. 34), give

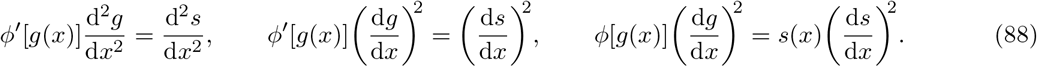

Applying the first two equalities to Eq. 86 produces Eq. 8 of the Results section.

Now we reintroduce noise *ζ* and assume it is independent across neurons and timesteps, with mean ⟨*ζ*⟩. If we average Eq. 84 over *ζ*, the numerator of the second term becomes

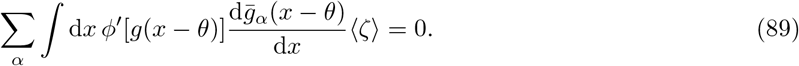

The integral vanishes because *g* is even and 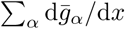 is odd. Thus,

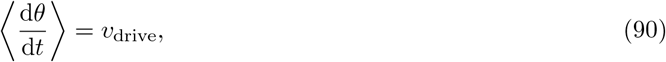

demonstrating that networks with independent noise still path integrate on average.

#### Diffusion *D*_input_ due to input noise

Independent noise *ζ* produces diffusion, a type of deviation in bump motion away from the average trajectory. It is quantified by the diffusion coefficient *D*:

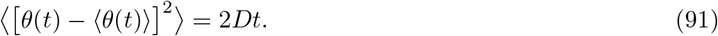

In terms of derivatives of *θ*,

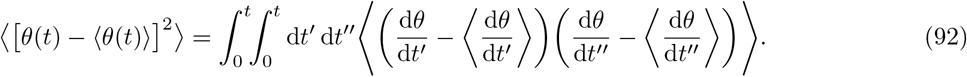

Equations 84 and 90 imply

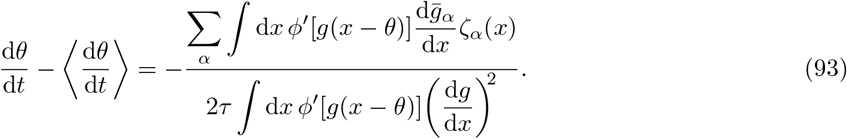

We then shift the dummy variable *x → x* + *θ*(*t*) and reintroduce explicit dependence on *t* to obtain

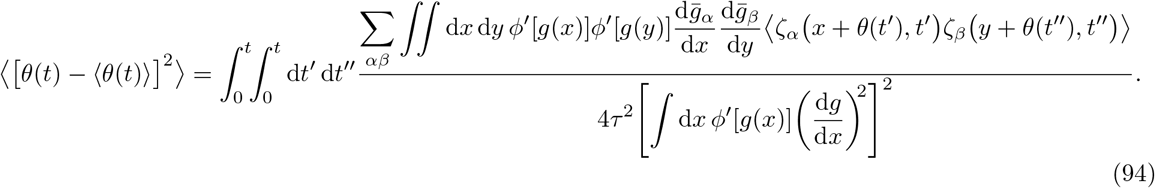

One class of independent *ζ* is Gaussian noise added to the total synaptic input, which represents neural fluctuations at short timescales. We assume it is independent across neurons and timesteps with zero mean and fixed variance *σ*^2^:

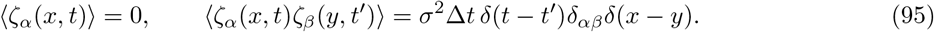

Δ*t* is the simulation timestep, which defines the rate at which the random noise variable is resampled.

Equation 94 then becomes, with the help of Eq. 78,

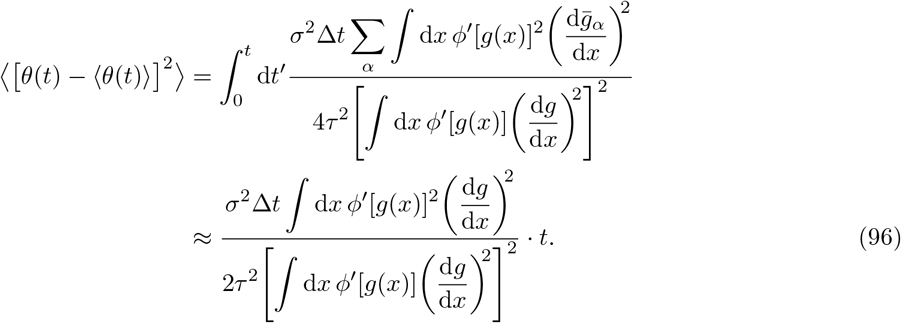

Reconciling this with the definition of the diffusion coefficient *D* in Eq. 91 yields

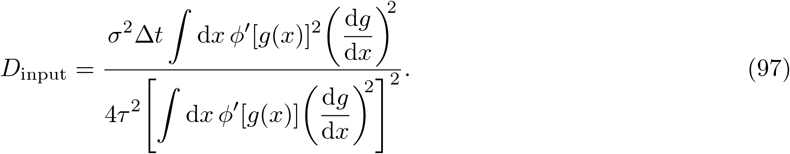

Applying Eq. 88 for a ReLU *ϕ* gives Eq. 10 of the Results section.

#### Diffusion *D*_spike_ due to spiking noise

Instead of input noise, we consider independent noise arising from spiking neurons. In this case, the stochastic firing rate *s* is no longer the deterministic expression in Eq. 34. Instead,

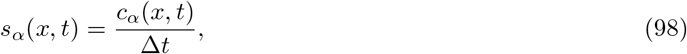

where *c* is the number of spikes emitted in a simulation timestep of length Δ*t*. We model each *c*_*α*_(*x, t*) as an independent Poisson-like random variable driven by the deterministic firing rate *ϕ*[*g*_*α*_(*x, t*)] with Fano factor *F*. It has mean *ϕ*[*g*_*α*_(*x, t*)]Δ*t* and variance *Fφ*[*g*_*α*_(*x, t*)]Δ*t*. Therefore,

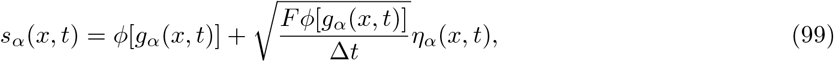

where each *η*_*α*_(*x, t*) is an independent random variable with zero mean and unit variance:

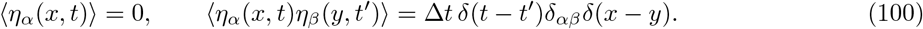

As in Eq. 95, the simulation timestep Δ*t* defines the rate at which *η* is resampled. By substituting Eq. 99 into Eq. 33, we see that spiking neurons can be described by deterministic firing rate dynamics with the stochastic noise term

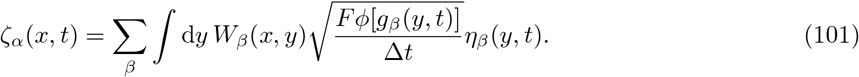

Now we calculate the diffusion coefficient produced by this noise. Equation 93 becomes

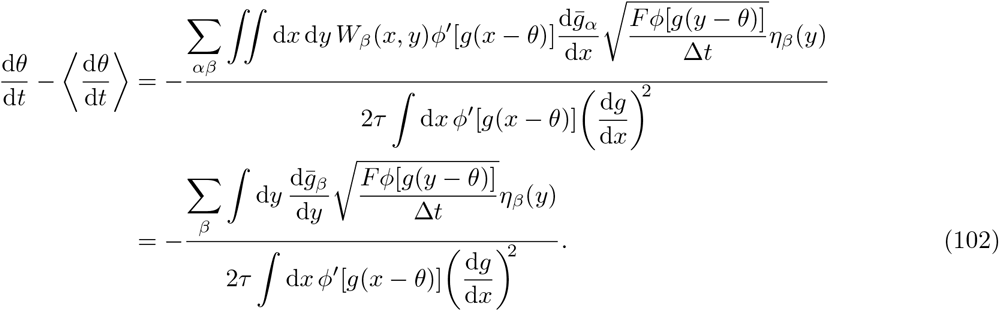

We used Eq. 79 to obtain the second equality. We then proceed as for input noise to calculate

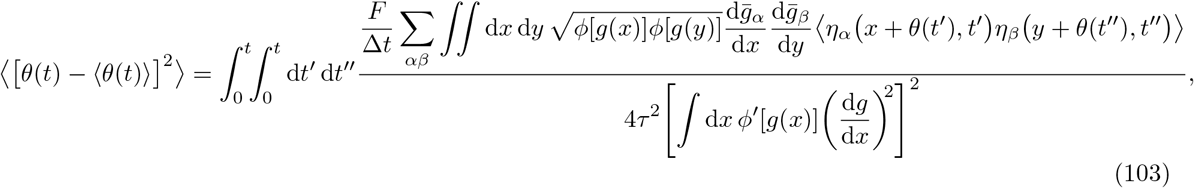

which yields the diffusion coefficient

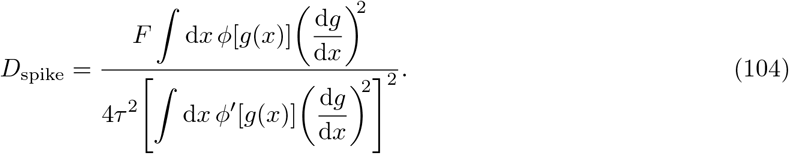

After applying Eq. 88 for a ReLU *ϕ* and setting *F* = 1 for Poisson spiking, we obtain Eq. 20 of the Results section.

#### Drift velocity *v*_conn_(*θ*) due to quenched connectivity noise

Suppose that we perturb the symmetric, translation-invariant *W* by a small component *V* representing deviations away from an ideal attractor architecture:

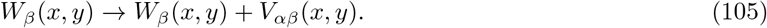

By Eq. 33, this produces the noise term

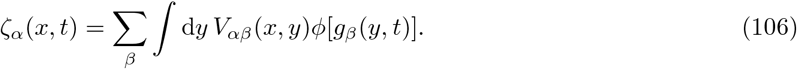

In contrast to input and spiking noise, this noise is correlated across neurons and time, so it cannot be averaged away as in Eqs. 89 and 90. Substituting Eq. 106 into Eq. 84, we obtain

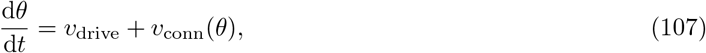

where the drift velocity is

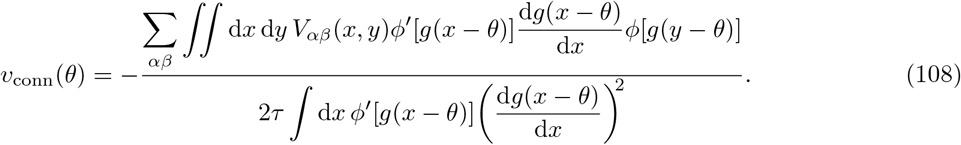

We have made the dependence on bump position *θ* explicit to illustrate how it influences *v*_conn_(*θ*). After applying Eq. 88 for a ReLU *ϕ*, we obtain Eq. 24 of the Results section.

We now make scaling arguments for speed difference (Eq. 30), speed variability (Eq. 31), and escape drive *b*_0_ (Eq. 26). To do so, we impose a ReLU *ϕ* and return to discrete variables to be explicit:

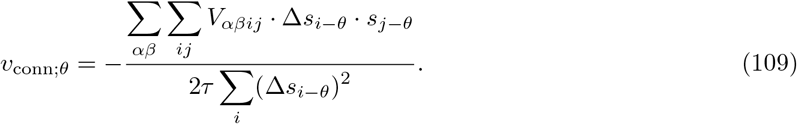

We need to understand how the numerator scales with *M* and *N*. It is a weighted sum of 4*N* ^2^ independent Gaussian random variables *V*_*αβij*_ and is thus a Gaussian random variable itself. It has zero mean, but its variance is proportional to *N* ^2^ *M* ^2^*/N* ^2^. The *N* ^2^ comes from the number of terms in the sum and the *M* ^2^*/N* ^2^ comes from the scaling of d*s/*d*x* (Eq. 11). In combination with the scaling of the denominator, we conclude that *v*_conn;*θ*_ is a Gaussian random variable with

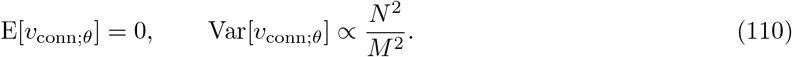

Equation 109 implies that *v*_conn;*θ*_ is correlated over *θ*. The weights for the sum over *V*_*αβij*_ are the firing rates and their derivatives for a bump centered at *θ*. If *θ* is slightly changed, almost the same entries of *V* will be summed over with similar weights. The amount of correlation across *θ* is determined by the degree of overlap in weights, and therefore, by the width and number of bumps. Let’s consider the effects of changing *N* and *M* on the covariance matrix Cov[*v*_conn;*θ*_, *v*_conn;*θ*_]. A larger *N* increases the bump width and the correlation length proportionally, so values of the main diagonal decay proportionally more slowly into the off diagonals. A larger *M* redistributes values among the diagonals by decreasing the bump width and adding more bumps, but it does not change the total amount of correlation. Thus,

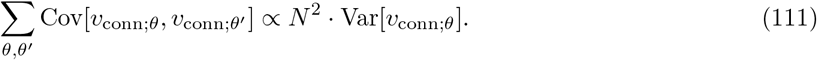

This allows us to evaluate

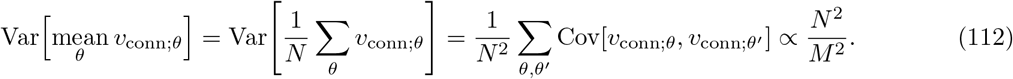

As a sum of zero-mean Gaussian random variables, mean_*θ*_ *v*_conn;*θ*_ is also a zero-mean Gaussian random variable. That means |mean_*θ*_ *v*_conn;*θ*_| follows a folded normal distribution, which obeys

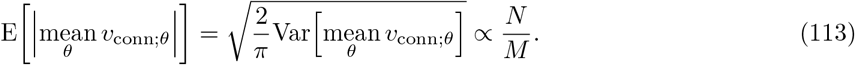

Combining this with Eqs. 12 and 14 produces the scalings for speed difference in Eq. 32. We now study speed variability, which involves the expression

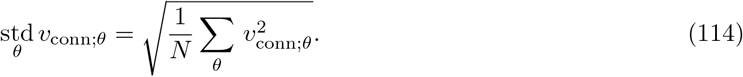

Since each *v*_conn;*θ*_ is Gaussian, the sum of their squares follows a generalized chi-square distribution. Its mean is the trace of the covariance matrix Cov[*v*_conn;*θ*_, *v*_conn;*θ*_], which is equal to *N* times the variance. Thus, by Eq. 110,

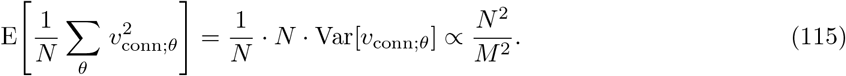

We are interested in the square root of the random variable on the left-hand side, and we anticipate its expected value to scale as the square root of the right-hand side. We can make this argument precise. Suppose *H* is a random variable with a probability distribution function *p*(*h*) that scales with a power of the parameter *B*. We can write

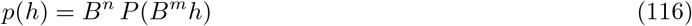

for exponents *n* and *m*, where the rescaled probability distribution function *P* does not scale with *B*. Conservation of total probability implies

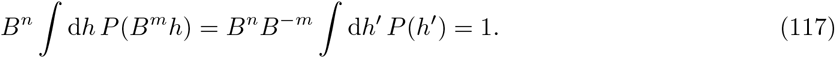

Thus, *m* = *n*. Next, suppose we know that E[*H*] *∝ B*^*o*^:

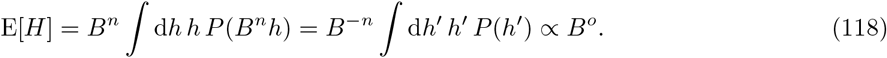

Thus, *n* = *−o*. We can now conclude that 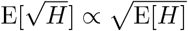

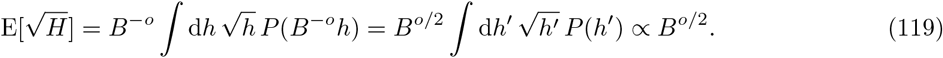

Applying this result to Eq. 115, we obtain

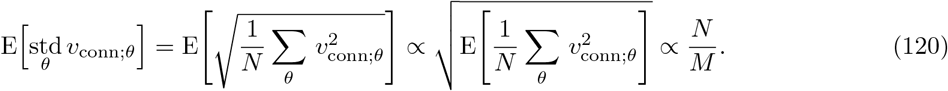

Combining this with Eqs. 12 and 14 produces the scalings for speed variability in Eq. 32.

The escape drive *b*_0_ involves the expression max _*θ*_|*v*_conn;*θ*_|. Extreme value statistics for correlated random variables is generally poorly understood. We follow Majumdar et al. (2020) and provide a heuristic argument for its scaling. We can partition *v*_conn;*θ*_ across *θ* into groups that are largely independent from one another based on its correlation structure. As discussed above, *v*_conn;*θ*_ is a weighted sum of independent Gaussian random variables *V*_*αβij*_ (Eq. 109). The weights are products between the firing rates *s*_*j−θ*_ and their derivatives Δ*s*_*i−θ*_ for a configuration centered at position *θ*. If we choose two *θ*’s such that bumps do not overlap, the corresponding *v*_conn;*θ*_’s will sum over different *V*_*αβij*_’s and will be independent. Thus, *λ/z* roughly sets the number of independent components, where *λ* is the bump distance and *z* is the bump width. This ratio does not change with *M* or *N* in our networks (Fig. 2F), so the maximum function does not change the scaling of |*v*_conn;*θ*_|:

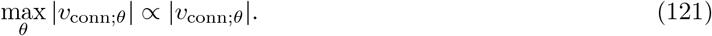

The scaling of E[|*v*_conn;*θ*_|] can be determined from Var[*v*_conn;*θ*_] through arguments similar to those made in Eqs. 116, 117, 118, and 119. Suppose we know that Var[*H*] *∝ B*^*o*^ and E[*H*] = 0. Then,

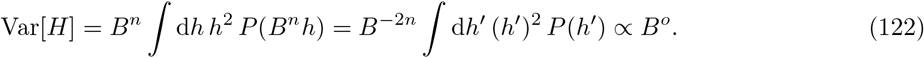

Thus, *n* = *−o/*2. We can now conclude that 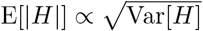:

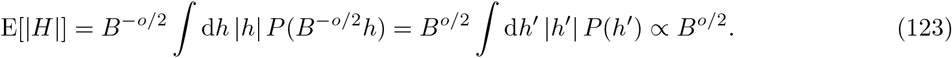

Applying this result to Eq. 121, we obtain

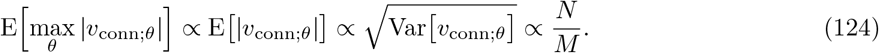

Combining this with Eqs. 12, 13, and 26 produces the scalings for the escape drive *b*_0_ in Eq. 27.

## Simulation methods

### Dynamics and parameter values

To simulate the dynamics Eq. 33, we discretize the network by replacing neural position *x* with index *i* and propagate forward in time with the simple Euler method:

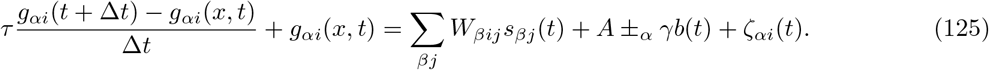

We use *τ* = 10ms. We use Δ*t* = 0.5ms and *A* = 1 for all simulations except those with spiking neurons. In the latter case, we use finer timesteps Δ*t* = 0.1ms and set *A* = 0.1ms^*−*1^. Synaptic inputs *g* and resting inputs *A* can be dimensionless for rate-based simulations, but they must have units of rate for spiking simulations. We use *γ* = 0.1 for rate-based simulations and *γ* = 0.01ms^*−*1^ for spiking simulations. In all cases, we run the simulation for 1000 timesteps before recording any data to form the bumps. To achieve the relationship in Eq. 13 for circular mapping, we rescale *γ* with network size *N* and bump number *M*:

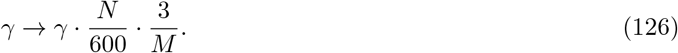

The connectivity *W* takes the form in Eq. 38. Unless otherwise specified, we use shift *ξ* = 2. To produce *M* bumps in a network of size *N*, we turn to Eq. 47 and set *l* = 0.44*N/M*. We use *w* = 8*M/N* ≈ 3.5*/l*. For the case of 2*l > N/*2, which corresponds to a one-bump network, the tails of the cosine function extend beyond the network size. Instead of truncating them, we wrap them around the ring:

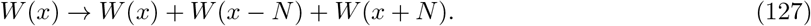

This procedure, along with the scaling of *w* with *N* and *M*, accomplishes Eq. 7 and keeps the total connectivity strength per neuron Σ_*i*_ *W*_*i*_ constant across all *N* and *M*, where *W*_*i*_ is the discrete form of *W* (*x*).

To generate the Poisson-like spike counts *c*_*αi*_(*t*) in Eq. 98, we rescale Poisson random variables:

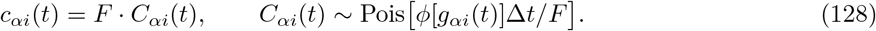

These counts will be multiples of the Fano factor *F*. To produce a *c*_*αi*_(*t*) whose domain is the natural numbers, one can follow Burak and Fiete (2009), who take multiple samples of *C*_*αi*_(*t*) during each timestep.

To obtain theoretical values in Figs. 3, 5, 7, and 8, we need to substitute the baseline inputs *g*_*i*_ into the appropriate equations. We use noiseless and driveless simulations to generate *g*_*i*_ instead of using Eq. 4.

### Bump position

We track the position *θ* of each bump using the firing rate summed across both populations *S*_*i*_(*t*) =Σ_*α*_ *ϕ*[*g*_*αi*_(*t*)]. We first estimate the positions of all the bumps by partitioning the network into segments of length ⌊*N/M*⌋. If *N/M* is not an integer, we skip one neuron between some segments to have them distributed as evenly as possible throughout the network. We sum *S*_*i*_(*t*) across all the segments and find the position *i*_0_ with maximum value. We perform a circular shift of the original *S*_*i*_(*t*) such that *i*_0_ is shifted to the middle of the first segment ⌊*N/*2*M*⌋. The purpose of this process is to approximately center each bump within a segment so that *S*_*i*_(*t*) drops to 0 before reaching segment boundaries. We then calculate the center of mass of *S*_*i*_(*t*) within each segment. After reversing the circular shift, these centers of masses are taken to be the bump positions.

As an alternative, we can obtain a bump position between 0 and *N/M* by simply computing the circular mean of *S*_*i*_(*t*) with periodicity *N/M*. However, this method does not track the position of each bump, so we do not use it.

### Path integration velocity and diffusion

To obtain our results in Figs. 3 and 5, we run each simulation for *T* = 5 s. To extract the bump velocity *v* produced by a constant drive *b*, we calculate the mean displacement Θ as a function of time offset *u*:

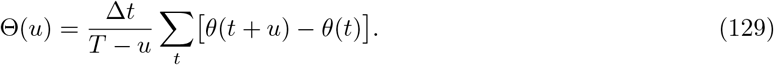

*θ* is the bump position. This equation averages over fiducial starting times *t*, which ranges from 0 to *T−u−*Δ*t* in increments of Δ*t*. We vary *u* between 0 and *T/*2 in increments of Δ*t*; the maximum is *T/*2 to ensure enough *t*’s for accurate averaging. We then fit Θ(*u*) to a line through the origin to obtain the velocity:

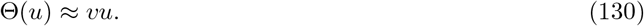

We calculate the diffusion coefficient *D* based on an ensemble of replicate simulations. In this section, angle brackets will indicate averaging over this ensemble. Following the definition of *D* in Eq. 92, we calculate each bump’s position relative to the mean motion of the ensemble:

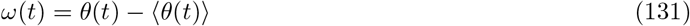

We compute squared displacements and then average over fiducial starting times to obtain a mean squared displacement for each bump as a function of time offset *u*:

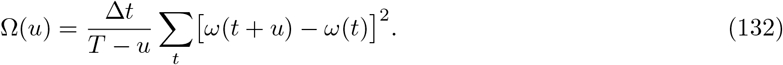

*t* and *u* span the same time ranges as they did for Θ. We average Ω(*u*) over the ensemble and fit it to a line through the origin to obtain the diffusion coefficient:

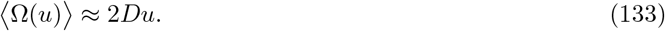

For simulations with *M* bumps, we arbitrarily assign identity numbers 1, …, *M* to bumps in each simulation. We perform ensemble averaging over bumps with the same identity numbers; that is, we only average over one bump per simulation. This way, we obtain separate values for each bump in Fig. 3E–H; nevertheless, these values lie on top of each other. In Fig. 3B, C, each point represents *v* averaged across bumps. To calculate the mean velocity ⟨*v*⟩ in Fig. 3E, F, we fit ⟨Θ(*u*)⟩ to a line through the origin. To estimate standard deviations for Fig. 3E–H and Fig. 5, we create 48 bootstrapped ensembles, each of which contains 48 replicate simulations sampled with replacement from the original ensemble. We calculate ⟨*v*⟩ or *D* for each bootstrapped ensemble and record the resulting standard deviation. In Fig. 5, each point represents *D* and its estimated standard deviation averaged across bumps.

### Trapping and position-dependent velocity

For simulations with connectivity noise, we determine the escape drive *b*0 (Fig. 7), the smallest drive that allows the bumps to travel through the entire network, by a binary search over *b*. We perform 8 rounds of search between the limits 0 and 1.28 and another 8 rounds between 0 and *−*1.28 to obtain *b*0 within an accuracy of 0.01. In each round, we run a simulation with the test *b* and see whether the bumps travel through the network or get trapped. Traveling through the network means that every position (rounded to the nearest integer) has been visited by a bump, and trapping means that the motion of at least one bump slows below a threshold for a length of time.

To obtain the position-dependent bump velocity *v*(*θ*) produced by connectivity noise when |*b*| *> b*0, we run a simulation until the bumps have traveled through the network. At each timestep, we record the positions of the bumps (binned to the nearest integer) and their instantaneous velocities with respect to the previous timestep. We smooth the velocities in time with a Gaussian kernel of width 10ms, which is the neural time constant *τ*. We calculate the mean and standard deviation of these smoothed velocities for each position bin.

### Mutual information

For simulations with input noise, we explore the mutual information between encoded coordinate and single-neuron activity (Fig. 6). To do so, we must generate data from which we can calculate *p*(*s*|*u*) in Eq. 22, for coordinate *u* ∈ *𝒰* and activity *s* ∈ *S*. We have chosen one set of conditions for performing this analysis, which we detail below.

We first choose to represent either a linear or circular coordinate, which we take to be position or orientation, respectively. We then choose to represent a narrow or wide coordinate range *u*_max_, which is 20 cm or 200 cm for position and 36 or 360 for orientation. We divide the range into 20 equally spaced coordinates such that 𝒰 = {*u*_max_*/*20,…, *u*_max_}. We convert these coordinates to network positions according to the mappings in Fig. 4. For each coordinate value *u*, we initialize 96 replicate simulations at the corresponding network position by applying additional synaptic input to the desired bump positions during bump formation. We run the simulations for 5 s, record the final firing rates, and bin them using 6 equally spaced bins from 0 to the 99th percentile across all neurons. All rates above the 99th percentile are also added to the 6th bin. These bins define the discrete 𝒮, and normalizing the bin counts produces *p*(*s*|*u*). We marginalize over *u* to obtain *p*(*s*), and *p*(*u*) is uniform. We can then use Eq. 22 to calculate the mutual information.

The 4 local cues in Fig. 6F–H correspond to 4 activity states 𝒮_cue_ separate from the 6 activity bins of the CAN neurons, 𝒮_neuron_. The joint sample space of a single neuron with cues is thus 𝒮 = 𝒮_neuron_*×* 𝒮_cue_ with 6*×*4 = 24 total states. We bin neural activity across these more numerous states, using the coordinate value *u* to determine the cue state value, to again calculate *p*(*s* |*u*) and then the mutual information.

We choose to calculate mutual information with single-neuron activities binned into 6 discrete states due to computational tractability. A better indication of encoding quality for the entire network would involve using the joint activity of multiple neurons. However, assuming the same binning process, that would involve estimating probability distributions over 6^*n*^ states for *n* neurons, which would require exponentially more replicate simulations per coordinate value than the 96 we use. Alternatively, one could reduce the dimensionality of the network activity by projecting it onto various attractor configurations, as done by Roudi and Treves (2008).

## Appendix

In this Appendix, we revisit many major results for input, spiking, and connectivity noise, but for either a different activation function *ϕ* (Fig. 9) or for connectivity strengths *W* that do not scale with bump number and network size (Fig. 10). To calculate theoretical predictions for each set of results, we need to substitute the baseline synaptic inputs *g* into the appropriate equations. They are obtained by running simulations without noise and drive. Notably, the theory still demonstrates close agreement with simulation results under these new conditions.

In Fig. 9, we use a logistic sigmoid activation function *ϕ* to convert synaptic inputs *g* to firing rates *s*:

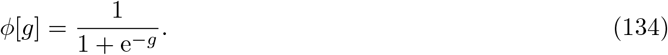

All results with this *ϕ* are qualitatively identical to those obtained with a ReLU *ϕ* in the Results section. To calculate theoretical values, we can no longer use equations from the Results, which are simplified for a ReLU *ϕ*. To calculate *D*_input,_*D*_spike,_*v*drive, and *v*_conn_(*θ*), we use Eqs. 97, 104, 86, and 108 from the Theoretical methods section instead.

In the Results section, we assumed that the connectivity strengths *W* obey Eq. 7 to maintain the same scaled bump shape across bump numbers *M* and network sizes *N*. In addition to the theoretical advantages of obtaining simple scaling relationships, this choice can be loosely biologically motivated. Consider the tuning curves of grid cells, which are thought to function as CANs. Their scaled shapes are roughly similar across modules (Stensola et al., 2012), which may differ in bump number (Gu et al., 2018; Kang and Balasubramanian, 2019; Khona et al., 2022), and across mammalian taxa from rodents to primates (Killian et al., 2012; Jacobs et al., 2013), whose brains certainly differ in neuron number. This crude observation supports the choice to maintain a fixed scaled bump shape across *M* and *N*. Nonetheless, in Fig. 10, we do not assume Eq. 7 and bump shape invariance. Instead, we fix *w* = 0.04 in Eq. 38, which fixes the maximum synaptic strength across all networks. This change produces qualitative differences only for circular mapping. Here, under circular mapping, networks with fewer bumps are more robust to all three forms of noise, and larger networks are more robust to connectivity noise. For the corresponding simulations in the Results section, no major dependence on bump number was observed.

## Acknowledgments

We are grateful to Steven Lee for sharing his code and to John Widloski for a careful reading of this manuscript.

